# Visualizing dynamic competence pili and DNA capture throughout the long axis of *Bacillus subtilis*

**DOI:** 10.1101/2023.05.26.542325

**Authors:** Jason D. Zuke, Rachel Erickson, Katherine R. Hummels, Briana M. Burton

**Affiliations:** Department of Bacteriology, University of Wisconsin - Madison; Microbiology Doctoral Training Program, University of Wisconsin - Madison; Department of Microbiology and Immunology, Harvard Medical School

**Keywords:** competence, natural transformation, pili, *B*. *subtilis*

## Abstract

The first step in the process of bacterial natural transformation is DNA capture. Although long-hypothesized based on genetics and functional experiments, the pilus structure responsible for initial DNA-binding had not yet been visualized for *Bacillus subtilis*. Here, we visualize functional competence pili in *Bacillus subtilis* using fluorophore-conjugated maleimide labeling in conjunction with epifluorescence microscopy. In strains that produce pilin monomers within ten-fold of wild type levels, the median length of detectable pili is 300nm. These pili are retractile and associate with DNA. Analysis of pilus distribution at the cell surface reveals that they are predominantly located along the long axis of the cell. The distribution is consistent with localization of proteins associated with subsequent transformation steps, DNA-binding and DNA translocation in the cytosol. These data suggest a distributed model for *B. subtilis* transformation machinery, in which initial steps of DNA capture occur throughout the long axis of the cell and subsequent steps may also occur away from the cell poles.

**Importance:** This work provides novel visual evidence for DNA translocation across the cell wall during *Bacillus subtilis* natural competence, an essential step in the natural transformation process. Our data demonstrate the existence of natural competence associated, retractile pili that can bind exogenous DNA. Furthermore, we show that pilus biogenesis occurs throughout the cell long axis. These data strongly support DNA translocation occurring all along the lateral cell wall during natural competence, wherein pili are produced, bind to free DNA in the extracellular space, and finally retract to pull the bound DNA through the gap in the cell wall created during pilus biogenesis.

## Introduction

Natural competence, the ability for bacteria to produce proteins that mediate the uptake of extracellular dsDNA, is widely distributed among the bacterial domain of life (1). DNA imported via natural competence is highly advantageous to cells and can be utilized in numerous, non-mutually exclusive ways. Once in the cytosol, the DNA can be metabolized to supply additional nucleotides and/or cellular carbon(2–4). If naturally competent cells encounter chromosomal DNA damage, the internalized DNA can be used for repair either through excision-mediated mechanisms or can be integrated into the resident chromosome via homologous recombination (5–8). Finally, the internalized DNA can confer novel genetic elements into the competent cell’s genome (9–11). The process wherein DNA is internalized and subsequently integrated into the competent cell’s chromosome is known as natural transformation and is one of the major drivers of horizontal gene transfer (HGT) in bacteria (12, 13).

The mechanisms governing Gram-positive natural transformation have been well characterized owing to work performed on model organisms such as *Streptococcus pneumoniae* and *Bacillus subtilis* (14). Transcription of genes involved in the production of DNA uptake and transformation proteins is activated through varied environmental cues (1). Once produced, a subset of these proteins must mediate the translocation of extracellular DNA across the negatively charged, 30-40 nm thick Gram-positive cell wall (15). The *comG* operon includes seven genes, each of which is homologous to components present in both type IV pilus (T4P) and type II secretion pseudopilus (T2SS) systems (16). These evolutionarily related systems employ a conserved set of proteins including at least one ATPase, at least one polytopic integral membrane protein, and a set of structural proteins known as pilins that work together to dynamically assemble membrane-bound, filamentous protein helices composed of repeating pilin subunits (17, 18). These filaments are known as either pili for T4P, or pseudopili for T2SS, with the main discriminating characteristic being filament length. T4P pili can be multiple microns long and readily observed in the extracellular space, while T2SS pseudopili are typically not long enough to be visible in the extracellular space (19). Crucially, both systems can produce filaments which span the cell envelope and reach the extracellular space (T2SS pilin genes must be overexpressed for this to occur (19)).

The *comG* operon mediates the production of seven proteins that work together to form a T4P filament, composed of hundreds to thousands of individual ComGC pilin subunits, that extends through the cell wall into the extracellular space (20–22). Once extended, the ComGC pilus binds to free dsDNA in the environment (20, 21). Dynamic retraction of the ComGC subunits back into the membrane shortens the pilus, transporting bound DNA through the gap formed in the cell wall, into contact with the membrane-bound DNA receptor/binding protein ComEA (23–26). After ComEA binding, DNA is brought into contact with the ComEC membrane channel (27, 28). The cytosolic protein ComFA aids in DNA entry across this channel via ATP hydrolysis (29–31). One strand of the incoming DNA is hydrolyzed, resulting in ssDNA in the cytosol (32–34). Through the action of various ssDNA binding proteins and the competence-specific RecA-loading protein DprA, RecA binds to the ssDNA and directs homologous recombination of the incoming ssDNA into the chromosome to complete natural transformation (35, 36).

Although this mechanism has been thoroughly researched, an interesting discrepancy remains regarding DNA translocation across the cell wall. While *S. pneumoniae* undoubtedly produces type IV pili to mediate initial DNA reception and translocation across the cell wall, all attempts to identify such a structure in *B. subtilis* have yielded negative results, despite *B. subtilis* carrying a *comG* operon that is both essential for natural transformation and is homologous to the *S. pneumoniae comG* operon (37, 38). Biochemical data indicate that *B. subtilis* produces multimeric ComGC structures associated with the cell wall (22, 39). However, no microscopy data or structural studies have confirmed the existence of *bona fide* pili or pseudopili, so the exact nature of *B. subtilis* translocation of transforming DNA across the cell wall is unclear.

In this investigation, we report that ectopic expression of select *comGC* cysteine substitution alleles (*comGC^Cys^*) in parallel with *comGC^WT^* allows for natural transformation to occur in *B. subtilis*. Expressing *comGC^Cys^* results in extracellular filaments that are labelable with a fluorescent maleimide-dye conjugate. We interpret these filaments to be pili composed mainly of ComGC. The pili can retract back towards the cell body and can bind to extracellular DNA. These data all support a model of DNA translocation across naturally competent *B. subtilis* cell walls by binding of DNA to pili, followed by pili retracting to transport bound DNA across the cell wall. We also find the localization of pilus biogenesis and DNA binding strongly biased to the periphery of the cell long axis. In addition, GFP-ComEA, and GFP-ComFA also reside predominantly along the long axis of the cell. Together the data suggest that DNA translocation across the cell wall is biased away from the poles, and even later steps in transformation may not be localized exclusively to or near the cell poles as has been previously reported (40, 41).

## Results

### ComGC cysteine substitution variants support transformation

Prior studies of natural competence associated pili, including the type IV competence pili of *V. cholerae* and *S. pneumoniae*, produced fluorescently labeled pili by using cysteine substitution variants of major structural pilins (20, 42). Since *B. subtilis* competence pilus biogenesis likely involves conversion of an intramolecular disulfide to an intermolecular disulfide between ComGC pilin subunits, we first assessed how the introduction of cysteine substitution variants affected transformability (22, 43). When *comGC^Cys^* alleles were ectopically expressed as the sole copy of *comGC*, transformation efficiency dramatically decreased by multiple orders of magnitude compared to wild type (Fig. S1). When *comGC^WT^* was expressed in the same strain background from the ectopic locus, however, a similar decrease in transformation efficiency was observed (Fig. S1E). This indicated to us that our ectopic expression construct had an intrinsic flaw that prevented proper complementation of the endogenous *comGC* deletion present in each strain. Importantly, this also implied that any strain ectopically expressing *comGC^Cys^* that maintained transformation efficiency near that of the isogenic strain ectopically expressing *comGC^WT^* was likely completely or mostly functional. Two alleles, *comGC^E56C^* and *comGC^S65C^*, allowed for transformation efficiency near that of *comGC^WT^*, therefore we decided to focus on *comGC^E56C^* and *comGC^S65C^*going forward (Fig. S1CD).

To prevent any problems arising from deletion of endogenous *comGC*, we simply opted to ectopically express *comGC^E56C^* or *comGC^S65C^* without altering the native *comG* operon, resulting in merodiploid strains expressing both *comGC^WT^* and *comGC^E56C^* or *comGC^S65C^*. We surmised that any potential pilus would be able to incorporate both ComGC^WT^ and ComGC^Cys^, allowing for pilus labeling via ComGC^E56C^ or ComGC^S65C^ subunits. These strains also carried an inducible *comK* allele to increase the fraction of the population that would express late competence gene products and thus facilitate microscopic analysis (44). Both *comGC^E56C^* and *comGC^S65C^* had limited negative effects on transformation efficiency when co-expressed with native *comGC*. The strains expressing *comGC^E56C^*and *comGC^S65C^* under control of the native *comG* promoter were cultured until one hour from maximum competence, at which point *comK* expression was induced. Donor DNA carrying a spectinomycin resistance gene at a separate ectopic locus was transformed after one hour of induction. The merodiploid strains producing the cysteine variants had transformation efficiencies within an order of magnitude of the matched wild type as well as a strain expressing only the endogenous copy of *comGC* (Fig. 1A). These transformation efficiencies were consistent with a functional apparatus for DNA translocation across the cell wall, which further encouraged us to continue using *comGC^E56C^*and *comGC^S65C^* in future experiments.

**Figure 1:**
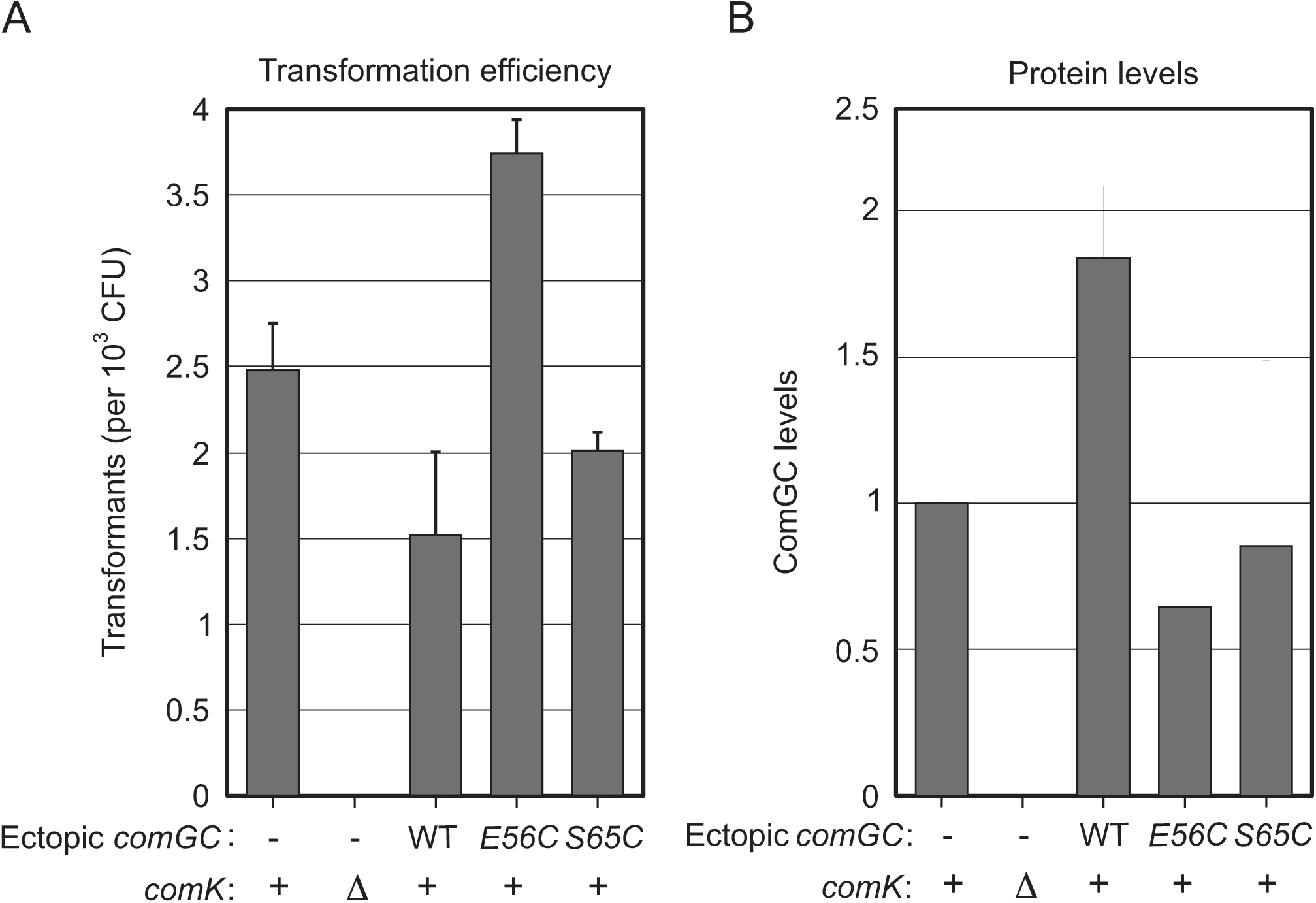
*B. subtilis* strains ectopically expressing *comGC^Cys^* are transformable. (A) Transformation efficiency of *B. subtilis* strains expressing *comGC^Cys^* from an ectopic locus. All strains carry inducible *P_xyl_-rbs-comK*. The presence and identity of the second *comGC* copy is indicated. The Δ*comK* strain did not have inducer added. The ratio of spectinomycin resistant transformants to total colony forming units was determined. The bar chart represents three biological replicates (error bars represent population standard deviation). (B) ComGC protein levels from whole cell lysates of each strain referenced in Fig. 1A. Cells from each strain were harvested after one hour of competence induction, total protein for loading normalization was visualized using Ponceau staining, and ComGC levels assessed through Western blotting using anti-ComGC 1° antisera. Normalized ComGC levels were calculated via ImageJ, using total protein as a loading control. Relative ComGC levels are shown relative to *P_xyl_-rbs-comK* and representative of samples taken from three biological replicates.

Next, we assessed ComGC pilin levels in the merodiploid strains, as well as an isogenic control strain lacking an ectopic *comGC* copy. Previous investigations into type II secretion system pseudopili have demonstrated a direct relationship between pilin levels and pilus length (45). Therefore, we sought to verify that any pili observed would be biologically relevant, rather than artifacts caused by increased levels of pilin present in *comGC^Cys^* mutants. Whole cell lysates of each strain were produced after inducing competence for one hour as described above and prepared for immunoblots with antiserum to detect ComGC (Fig. 1B). No ComGC was detected for an uninduced Δ*comK* culture, confirming ComGC-specific signal. ComGC levels for both strains carrying *comGC^Cys^*at *lacA* were within ten-fold of the isogenic control strain carrying only the endogenous *comGC* copy (Fig. S2). We note that although there was much higher variability in ComGC levels in the merodiploid strains expressing *comGC^E56C^* or *comGC^S65C^* compared to strains with either a single or two copies of wild type *comGC,* these measurements support that any extended pilus identified is not simply due to excess ComGC subunits in the cell but is physiologically relevant.

### Multiple ComGC cysteine substitution variants produce extracellular filaments

With multiple transformable *comGC^Cys^* alleles identified, we next wanted to determine if strains expressing either allele produced pili observable in real-time via epifluorescence microscopy. To this end, we employed the widely used maleimide labeling method that has successfully identified natural competence-associated type IV pili from both Gram-negative and Gram-positive species (20, 42). This method utilizes the highly selective reactivity of maleimide towards thiol moieties. If a pilus is produced, and the major constituent pilin subunit contains an unpaired cysteine that faces the solvent, maleimide conjugates in the solvent will covalently bond with the free thiol group on each available ComGC^Cys^ cysteine residue. If the maleimide is conjugated to a fluorophore, epifluorescence microscopy can be used to identify fluorescent pili and observe their dynamic production and depolymerization in real-time.

To perform maleimide labeling, we first cultured *comGC^WT^*, *comGC^E56C^*, and *comGC^S65C^* to competence as described above. Samples of each culture were incubated with Alexa Fluor 488 C_5_ Maleimide in the dark, washed, and deposited on agar pads for microscopic analysis. Both *comGC^E56C^*and *comGC^S65C^* produced stubby, filamentous structures emanating from the cell periphery that resembled short pili (Fig. 2A). Critically, there was not a single instance of such structures being produced by *comGC^WT^* treated with AF488-Mal (Fig. 2A and Fig. S3). This demonstrates that observation of these filaments depends on the presence of the ComGC^Cys^ monomers, lending credence to the idea that ComGC is incorporated into filaments that project into the extracellular space. Due to this finding, and the wealth of information indicating ComGC’s homology to other known pilins in terms of both form and function, we will henceforth refer to these filamentous structures as “ComGC pili”.

**Figure 2:**
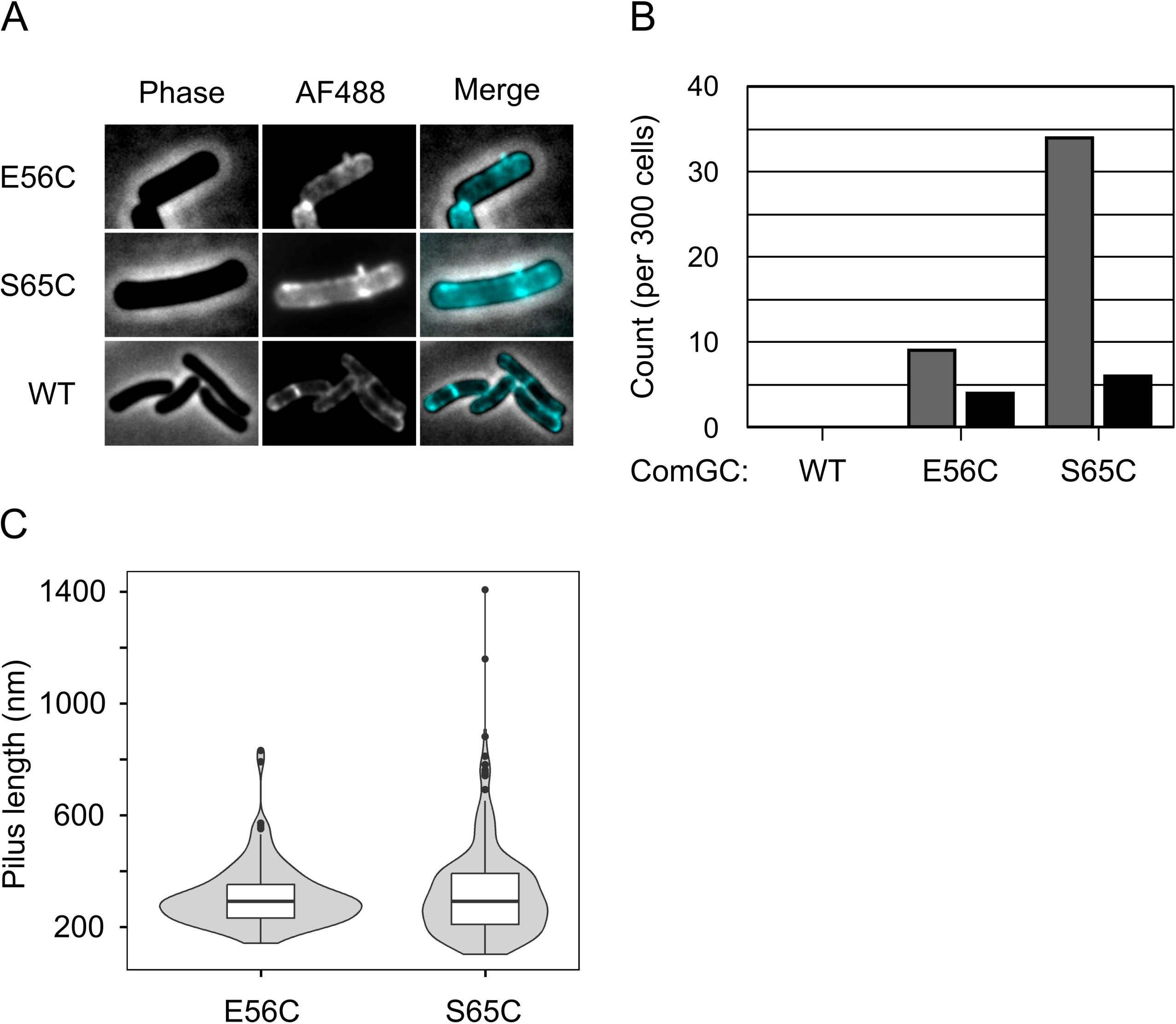
Strains expressing *comGC^Cys^* variants produce extracellular filaments composed of ComGC^Cys^ monomers. (A) Representative images of *B. subtilis* strains producing ComGC^Cys^ extracellular filaments. Competence was induced for one hour, followed by labeling with 25 µg/mL Alexa Fluor 488-maleimide for 20 minutes. Labeled cells were deposited on agar pads and imaged. (B) Quantification of ComGC pilus production in the same strains and culturing conditions referenced in Fig. 2A. The number of pili (black bars) or pilus-like elements (grey bars) was determined. External foci were defined as any discrete region of fluorescence above background that occurred within 0.5 µm of a cell, whereas pili were defined as any filamentous region of fluorescence contiguous to the boundary of a cell. Quantification was performed on 300 individual cells from each culture. (C) Quantification of ComGC ^E56C^ and ComGC ^S65C^ pilus lengths. Violin plots of 133 ComGC ^E56C^ pili and 233 ComGC ^S65C^ pili measured from cell periphery to tip of pilus. Mean lengths are 310 and 320 nm respectively.

The pilus production capacity of both strains was assessed quantitatively (Fig. 2B). Both intact pili that remained co-localized with the cell body, as well as regions of bright fluorescence near the periphery of the cell that we surmised were sheared pilus fragments (see discussion), were identified for 300 individual cells of each strain. While both strains produced both classes of signal, *comGC^S65C^* consistently produced a greater number of pili and pilus-like features compared to *comGC^E56C^*. We decided to move forward in our analysis using *comGC^S65C^* based on these results. Qualitatively, these ComGC pili were notably shorter than competence pili analyzed for other species across the bacterial domain (20, 42). Therefore, we assessed the distribution of pilus lengths for comparison (Fig. 2C). The pili produced by both *comGC^E56C^* and *comGC^S65C^*strains were indeed quite short, with mean pilus lengths of 0.32 µm and 0.33 µm and standard deviations of 0.11 µm and 0.17 µm, respectively. The implications of this size distribution will be expanded upon in the discussion.

### ComGC pili bind to DNA

One of the outstanding questions for *B. subtilis* DNA internalization is the mechanism by which transforming DNA is translocated across the cell wall (14). The prevailing hypothesis, that a pilus-like structure is formed which binds DNA and retracts to convey DNA to the membrane-localized DNA receptor ComEA, can now be tested directly with the aid of ComGC pilus maleimide labeling (14). First, we sought to address whether binding to DNA by ComGC pili occurs in the extracellular space during natural competence. To this end, an ∼4.5 kb PCR fragment from the *ycgO* locus was produced and covalently labeled with Cy5 fluorophores along the length of DNA. This labeled PCR product can be co-incubated with maleimide labeled cells to probe interactions between ComGC pili and extracellular DNA in real-time via time lapse epifluorescence microscopy (20, 42).

After silica column purification of the labeled PCR product, *comGC^S65C^*was maleimide labeled as described previously. To ensure that we could visualize all possible pili around the periphery of the cell, we collected Z-stacks with a small step size (0.2 µm). Collecting the z-stacks also generated a time lapse of the cells with roughly five second intervals between images. In numerous instances labeled PCR product was already co-localized with pili at the onset of imaging, and in other cases migrated towards pili and subsequently remained co-localized with the pili (Fig. 3 and Movie S1). Even subtle movements of pili were mirrored by co-localized PCR product over time. The persistent binding of PCR product through time, the coordinated movement of co-localized PCR product and pili, as well as the real-time migration and co-localization of PCR product to pili, would be highly unlikely to occur unless a *bona fide* binding interaction was being formed between pili and PCR product. These data provide direct evidence of the binding of DNA to *B. subtilis* ComGC pili.

**Figure 3:**
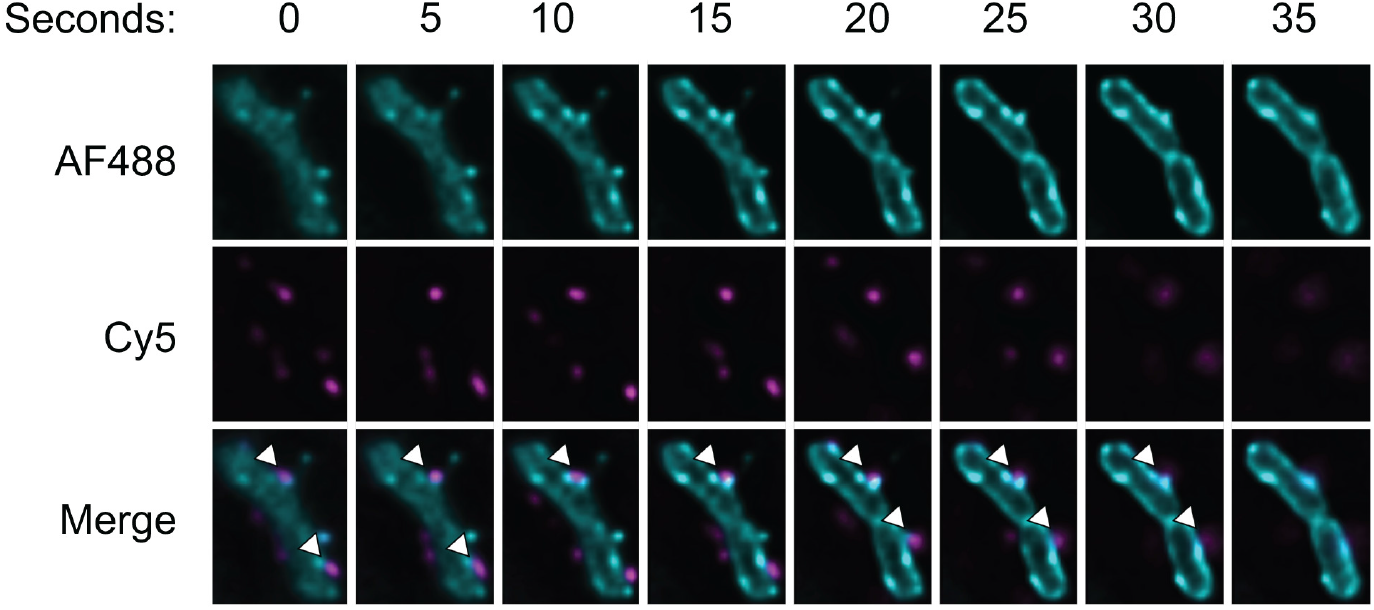
ComGC pili bind to extracellular DNA. Representative time series of ComGC pili binding to fluorescent DNA. ComGC^S65C^ pili were labeled as described previously, and fluorescently labeled PCR product was added to the cells prior to deposition on agar pads and imaging. Arrows indicate co-localizing pili and DNA.

### ComGC pili are retractile

Once a ComGC pilus has formed and bound to extracellular DNA, a translocation step must ensue to move the bound DNA across the cell wall, where it can then presumably interact with the membrane-bound DNA receptor ComEA (24). Given that type IV pili are dynamic structures that can retract back into the membrane by the sequential disassembly of pilin monomers into the membrane, the most parsimonious mechanism of translocating DNA across the cell wall is simply depolymerizing the ComGC pilus (17). Maleimide labeling of ComGC pili allows us the opportunity to study the dynamics of pilus production and depolymerization, which allows us to verify whether DNA translocation is driven by ComGC pilus retraction.

To assess ComGC pilus retraction, *comGC^S65C^* was maleimide labeled and deposited on agar pads as described previously. Time lapse imaging was performed using ten second intervals for one to two minutes to search for pili in the process of retraction (Fig. 4A and Movie S2). ComGC pilus retraction events were documented, wherein the length of a pilus would steadily decrease until only a strong focus of signal would persist at the cell periphery. For each pilus retraction event documented, the difference in pilus length between sequential frames was determined (44 total intervals across 10 pili). Interestingly, we observed two populations of retractile pili, one being considerably faster than the other. The median retraction rate for the “slow” group was 3 nm/sec, while the “fast” group was 12 nm/sec. These retraction rates were both notably slower than those calculated for competence pili of either *V. cholerae* or *S. pneumoniae*, which are much more in line with other observed type IV pilus systems (20, 42). All members within the fast population did not have statistically significant differences in average retraction rate nor did all members within the slow population (ANOVA, p = 0.36 and p = 0.69 respectively). However, a statistically significant difference was observed when the entire fast population and the entire slow population set of retraction rates were compared (T-Test, p = 3.4 x 10^-7^). These data will be examined further in the discussion section.

**Figure 4:**
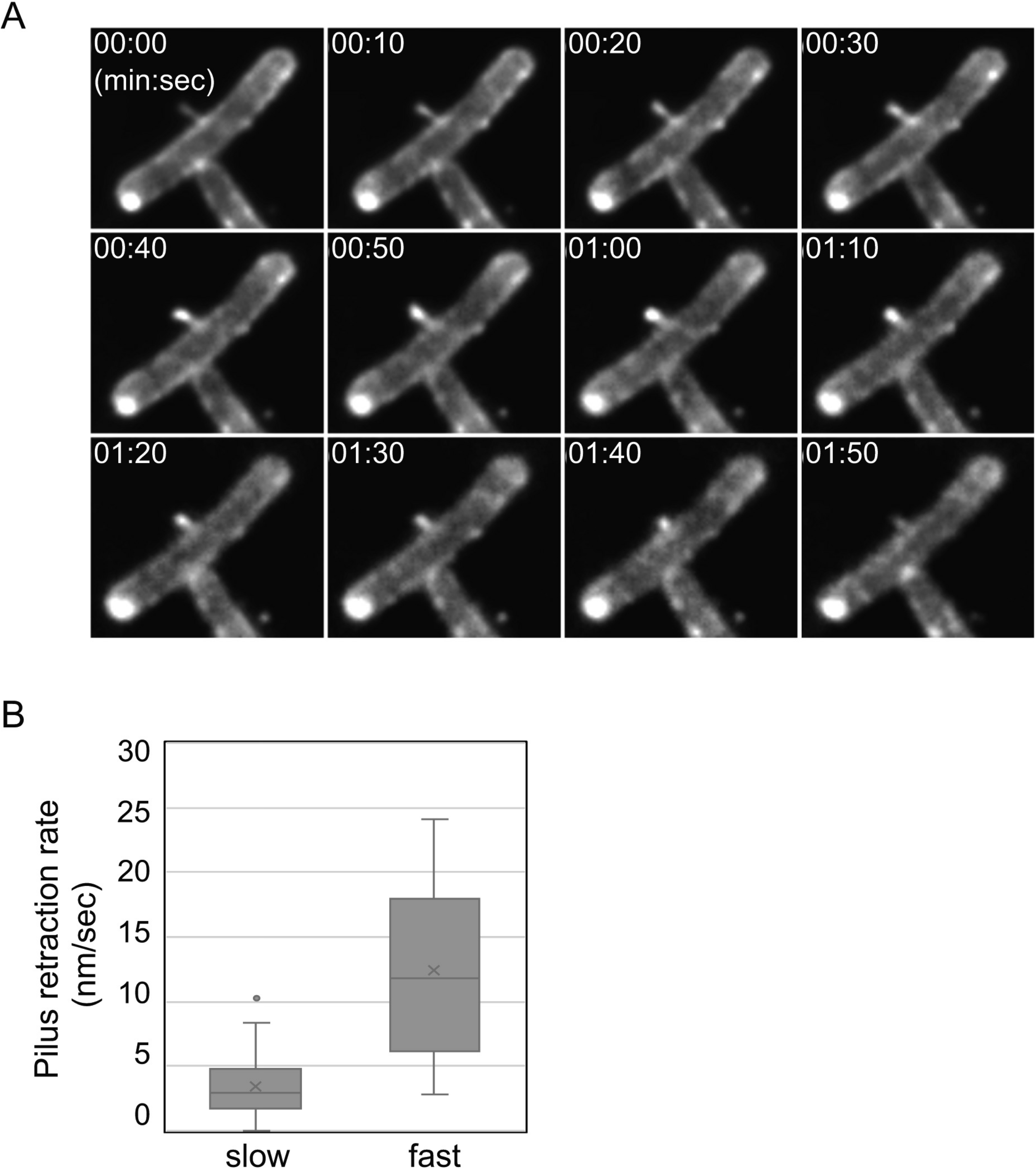
ComGC pili are retractile. (A) Representative time series of ComGC pilus retraction. *comGC^S65C^* was induced, labeled with Alexa Fluor 488-maleimide, and imaged on an agar pad as described previously. Cells were imaged every 5 seconds. (B) Box-and-whisker plots of ComGC^S65C^ pilus retraction rates drawn from the “slow” population (n = 4 pili) and the “fast” population (n = 6 pili). Individual retraction intervals from data like the example shown in (A) were combined and plotted (n = 22 intervals from each population).

### ComGC pili, DNA, ComEA, and ComFA localize along the long axis of competent cells

Our ability to identify sites of pilus biogenesis presents the opportunity to reassess the spatiotemporal dynamics of DNA uptake during natural competence. DNA uptake has been previously shown to occur primarily at or near the cell poles, perhaps taking advantage of a natural structural weakness in the cell wall region at the interface of the lateral and polar cell wall sections (40, 41). To probe this hypothesis, we analyzed the localization of ComGC pilus biogenesis in *comGC^S65C^*, as well as the localization of stably bound fluorescent PCR product. If DNA uptake occurs predominantly at or near the cell poles, we should expect the majority of assembled pili, and the majority of bound DNA, to localize close to the cell poles during natural competence.

The analysis of ComGC pilus localization, employing approximately 230 individual pilus biogenesis events, demonstrated that pili emanate along the long axis of competent cells at the cell periphery, with relatively few events occurring at or near the cell poles (Fig. 5A). In agreement with these data, fluorescent DNA that was stably bound to pili or the cell periphery primarily localized along the long axis of competent cells (Fig. 5B). To further assess this model of DNA uptake, we compared the distribution of ComGC pili to subcellular localization of functional GFP-ComEA and GFP-ComFA. Since ComEA serves the vital role of binding incoming DNA at the cell membrane, the protein should localize at the sites of initial DNA entry, which presumably coincides with the location of pili biogenesis. In accordance with this hypothesis, GFP-ComEA localizes along the long axis of competent cells, just like ComGC pili and bound DNA (Fig. 5C). There are two possibilities for the next phase of DNA entry. The handoff from ComEA to the ComEC channel could occur near the site of pilus biogenesis, or it could occur at a secondary location. ComFA, which powers DNA translocation across the cell membrane, thus should serve as a proxy for the cytoplasmic internalization step. Indeed, GFP-ComFA, like all the other competence proteins and DNA assessed above, localizes near the cell periphery along the long axis of competent cells (Fig. 5D). It would appear, therefore, that DNA translocation across the cell wall, and entry into the cytoplasm, need not occur proximal to a cell pole.

**Figure 5:**
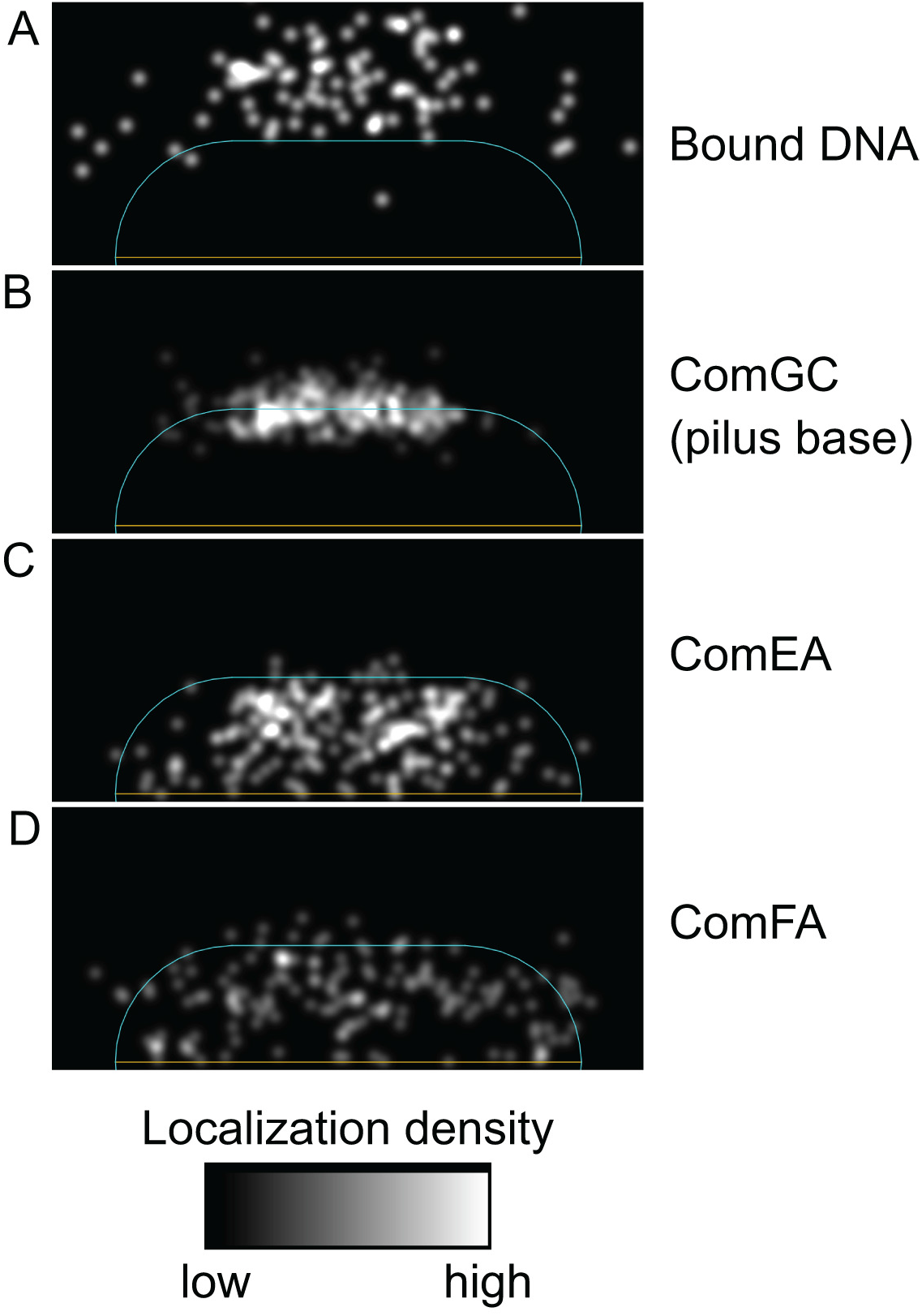
ComGC pili, DNA, ComEA, and ComFA localize along the long axis of competent cells. (A) Heatmap of *Label* IT-Cy5 (Mirus) treated 4.5 kb PCR product co-localizing with pili or discrete foci of ComGC^S65C^ fluorescence, along a normalized cell body (n = 93). Imaging was performed as described in Fig. 3, and localization analysis was performed using the MicrobeJ version 5.11x plugin for ImageJ. (B) Heatmap of *comGC^S65C^* pilus biogenesis locations along a normalized cell body (n = 232). (C) Heatmap of GFP-ComEA foci (n=200) along a normalized cell body. (D) Heatmap of GFP-ComFA foci (n-148) along a normalized cell body.

## Discussion

In *Bacillus subtilis*, the *comG* operon is essential for natural transformation (37). The ComG proteins of *B. subtilis* have been assumed to be involved in the production of a pilus capable of conveying extracellular DNA across the cell wall. This idea stems from multiple lines of evidence: each protein coded for in the operon has homology to components of either type IV pilus or type II secretion pseudopilus systems, there is biochemical evidence supporting multimerization of ComGC associated with the cell wall, and more recently ComGC pili have been demonstrated in *Streptococcus pneumoniae* (16, 20, 22, 39). However, direct evidence of ComG-mediated pilus production in *B. subtilis* has been lacking, with no microscopy or structural studies supporting pilus biogenesis. Here we demonstrate that *B. subtilis* in the naturally competent state produces ComGC-based pili. These pili are capable of stable binding of DNA, and they can retract dynamically back towards the cell membrane after assembly. ComGC pili were also found to localize along the long axis of the cell at the cell periphery, mirroring the localization of bound DNA and the downstream essential competence proteins ComEA and ComFA we observed.

We demonstrated that *comGC^E56C^*and *comGC^S65C^* alleles were functional by complementation of *ΔcomGC* for transformation efficiency (Fig. S1). These *comGC^Cys^*variants complemented almost as well as *comGC^WT^*, further supporting their functionality. The *comGC* complementation construct itself caused massive decreases in transformability regardless of the *comGC* allele used, and no pili were ever observed in either of the *ΔcomGC* complementation strains (Fig. S4). The most likely explanation for the poor transformability of these strains is a polar effect due the introduction of a sub-promoter into the *comG* operon during native *comGC* deletion (46). There are four additional pilin gene homologues downstream of *comGC* in the operon (47). Overexpression of certain pilins can be inhibitory for pilus production, so it’s feasible that the sub-promoter could cause overexpression of *comGDEFG*, and at least one gene product could inhibit pilus biogenesis (48). Regardless of the exact reason, the failure of the complementation strains to produce pili spurred us to try another approach to produce biologically relevant, visualizable pili.

Rather than alter the endogenous *comG* promoter, we instead took a less perturbative approach and expressed *comGC^Cys^* alongside native *comGC*. In stark contrast to the *comGC* complementation strains, we demonstrated that *comGC^E56C^* and *comGC^S65C^*merodiploid strains closely approximated wild type transformation efficiencies (within two-fold, Fig. 1A). The levels of total ComGC in both *comGC^E56C^*and *comGC^S65C^* were variable and slightly lower than for the isogenic control (Fig. 1B). While this may suggest that the ComGC pili we observed were artifactually shortened due to a smaller ComGC pool, it must be noted that the ComGC peptide used to raise the ComGC antiserum we employed for Western blotting included both positions E56 and S65 (38). It is therefore possible that the ComGC antiserum may not recognize ComGC^E56C^ or ComGC^S65C^ as efficiently as ComGC^WT^ since the binding epitope could have been altered. Total ComGC levels in *comGC^E56C^* and *comGC^S65C^* may therefore be roughly equivalent to the isogenic control, making the pili we observe reflective of those found in wild type cells, or possibly slightly shorter.

The mean pilus length measured in *B. subtilis* of 0.33 µm with simultaneous expression of *comGC^WT^* and *comGC^S65C^*(Fig. 2C) is notably shorter than the mean lengths of competence pili identified in *V. cholerae* or *S. pneumoniae* of 1 µm and 0.5 µm respectively (20, 42). The somewhat smaller difference between *B. subtilis* and *S. pneumoniae* ComGC pili may be attributable to pilus measurement error. Exact pilus start and end points had to be manually assigned based on the phase-contrast and epifluorescence images taken, which could lead to length differences based on user interpretation. Additionally, we determined pilus start points to be at the border of the phase-contrast (cell body) signal, not the border of the epifluorescence (ComGC) signal. There was generally a gap between the internal ComGC signal and the external cell body signal, so pilus lengths would have been increased had pilus start points been marked at the ComGC signal border. Another likely source of variation could be the presence of endogenous cysteine residues in *B. subtilis* ComGC. An off-target, BdbDC-mediated disulfide bond between an endogenous cysteine on one ComGC monomer and the mutant cysteine of another ComGC^Cys^ monomer might result in early termination of pilus elongation due to conformational changes at the pilus base that inhibit proper coordination of the pilus biogenesis apparatus. *S. pneumoniae* ComGC, in contrast, contains no endogenous cysteine residues that could improperly form disulfide bonds with the ComGC^Cys^ cysteine (see PDB 5NCA), negating this possibility. And finally, the observed length difference may simply stem from intrinsic differences in the protein sequences of each ComG protein comprising the two systems.

The larger difference between *V. cholerae* and *B. subtilis* pilus lengths may stem from differences in how force for pilus biogenesis is generated. *V. cholerae* employs both a dedicated extension (PilB) and a retraction (PilT) ATPase, whereas *B. subtilis* only has one identifiable pilus ATPase homologue (ComGA) (16, 42). PilB ATPase activity may simply be faster and/or more processive than that of ComGA, which would grant PilB greater ability to incorporate new pilin subunits into a growing pilus prior to disassembly than ComGA, making the average pilus longer for PilB-polymerized pili. Further investigation of pilin levels and extension/retraction ATPase activities are warranted to produce a complete picture of how competence pilus lengths are established.

Our work provides evidence of direct *B. subtilis* ComGC pilus-DNA interactions with physiological levels of ComGC proteins (Fig. 3). Such interactions are consistent with data from a diverse set of naturally competent bacteria, including *V. cholerae*, *S. pneumoniae*, *Neisseria gonorrhoeae*, and *Thermus thermophilus* that demonstrate either direct binding of pili to DNA or individual pilins to DNA (20, 42, 49, 50). We also demonstrate that *B. subtilis* ComGC pili are retractile in nature, as has been observed for *V. cholerae* and *S. pneumoniae* competence pili (Fig. 4) (20, 42). Intriguingly, we observed two distinct populations of retracting pili with variable retraction rates: a slow population that retracts with a median rate of 3 nm/sec and a fast population that retracts with a median rate of 12 nm/sec (Fig. 4C). The slow population may be retracting spontaneously, whereas the fast population is most likely actively retracting via ComGA activity. Spontaneous retraction of *V. cholerae* competence pili has been observed with a notably slower retraction rate compared to active retraction, strengthening this idea (42, 51, 52). Moreover, the clear distinction in retraction rates between the two populations suggests that a specific process increases the retraction rate of the fast-retracting pili. The most parsimonious explanation is, of course, that the pilus ATPase homologue ComGA is promoting active retraction events within the fast population.

The retraction rate for the fastest ComGC pili observed (median = 12 nm/sec) is much slower than for *V. cholerae* (median ∼ 100 nm/sec) and *S. pneumoniae* (median ∼ 80 nm/sec). We consider two possible explanations for this difference. First, the rate differences could be reflective of differences in the enzymatic activities of the ATPases utilized for retraction across these systems (Fig. 4B) (17). Alternatively, the disulfide isomerization involved in assembly and disassembly of ComGC pili in *B. subtilis* may impact the retraction rate. Dissection of the contribution of the disulfide is not trivial, as disulfide bond formation is necessary for ComGC pilus biogenesis (22), but future mechanistic studies to address this question will be valuable.

Subcellular localization analysis of ComGC pili and of associated DNA produced a localization pattern quite distinct from the previously reported polar localization patterns of other late competence gene products, including ComGA and ComEC (40, 41). We found that ComGC pili and associated DNA both predominantly localize across the long axis of the cell (Fig. 5AB), with little clustering of ComGC pilus biogenesis. Additionally, ComEA and ComFA were seen to localize along the long axis of competent cells (Fig. 5CD). These localization patterns are consistent with the predominant distributions reported for DNA bound at the cell membrane (25). These observations suggest that the initial step of mobilizing extracellular DNA across the peptidoglycan layer for ComEA binding at the cell membrane may occur throughout the surface of the cell, predominantly along the long axis, and may even be excluded from the cell poles. One possible explanation for this is that during natural competence the lateral cell wall is mostly static, as the cells have ceased growth and elongation, and any remaining peptidoglycan remodeling is likely occurring at the cell poles at sites of cell separation (53–55). This may facilitate ComGC pilus formation at this region, since steric clashes with active cell wall remodeling systems could potentially be reduced, and any channels formed in the cell wall may be more likely to remain open for extended periods of time.

While our data for ComEA localization are largely consistent with previous reports, our analysis differs significantly for ComFA localization (25, 40). We observed GFP-ComFA puncta throughout the long axis of the cell (Fig. 5D), whereas polar ComFA-YFP localization had been observed previously. This difference is of critical importance since the localization of ComFA to the cell poles, as well as that of other late competence proteins such as ComEC and ComGA, led to the conclusion that DNA translocation across the cell membrane occurred at or near the cell poles (40). Our data, on the other hand, are consistent with DNA entry points distributed along the lateral cell membrane.

To address the disparities, we consider DNA uptake in the context of the two-stage model for transformation (56). First, our data may best represent the location of the first step of DNA uptake, where DNA crosses the cell well. The localization of pilus biogenesis and retractile pili that interact with DNA provides a direct view of active competence pili. In contrast, the polar localization of ComGA holds the implicit assumption that the foci are coincident with active protein. (40, 57). Given the necessity of ComGA for pilus assembly, functional ComGA should localize to the sites of pilus biogenesis (i.e. along the cell long axis), but that has not been observed (39). This raises questions as to the biological relevance of the observed localization patterns of the late competence protein fluorescent fusions used in previous studies. Subpopulations of proteins are often sufficient for biological activity, and active fractions do not always correspond to the visualized population (58). This could be the case for the prior results with ComGA fusions.

The location of the second stage of DNA import, across the cell membrane as assessed by ComEC and ComFA localization, remains less clear. Studies from multiple groups have come to the same conclusion regarding polar localization of ComFA (40, 57). The constructs used in the respective localization experiments differ from ours in placement of the fluorescent tag. The data presented here use a complementing amino-terminal fluorescent fusion to ComFA. Consistent with prior published work, we observe predominantly polar localization when the fluorescent protein is at the carboxy-terminal end of ComFA. However, there is significant proteolysis of the fluorescent tag and free ComFA in these cells, reducing confidence in these fusions as reporters of active protein. Thus, we favor the amino-terminal fusion presented here as reporting on functional localization. However, from these data alone we cannot rule out that DNA translocation across the cell membrane could occur at cell poles. In such a system, DNA that had been captured by ComEA after translocation across the cell wall would most likely be transported to the cell pole, where the DNA would then be internalized (59).

Critically, our results expand on the mechanism of DNA reception and translocation across the cell wall during Gram-positive natural competence, where models have thus far relied on observations made solely using *S. pneumoniae* as a model organism. Our observations are most consistent with the following model of DNA translocation across the cell wall. During natural competence, the protein products of the *comG* operon work together to generate an extracellular pilus that is comprised primarily of ComGC along the long axis of the cell. These pili are dynamic and retract stochastically. At some point, dsDNA in the environment binds to the pilus surface. At some point after DNA binding, the pilus will retract back into the membrane, which will consequently pull the bound DNA through the gap left in the cell wall. Once across the cell wall, the DNA will bind to ComEA at the cell membrane, and translocation into the cytoplasm will commence, either at a cell pole or along the long axis of the cell.

Numerous questions remain unresolved regarding the mechanism of DNA translocation across the cell wall. Are each of the 5 pilin homologues in the *comG* operon present in the competence pilus, and if so, where are they located? Each gene in the *comG* operon is essential for transformation, so presumably each pilin is involved to some degree with pilus biogenesis and/or function (37). In *S. pneumoniae*, it was recently discovered that ComGC, ComGF, and ComGG were present throughout pili, although no other pilins were detected (60). It is possible that the other pilins (ComGD and ComGE) are located at the tip of the pilus in single copy and are responsible for DNA binding. This is consistent with the role of low abundance “minor pilins” in initiation of pilus assembly and interactions with the environment (17, 60, 61). Identification of minor pilin point mutations that allow for DNA-binding-deficient pili to be produced, as was achieved for *V. cholera* competence pili, would support this hypothesis (42). With successful methods to visualize ComGC pili and DNA capture in *B. subtilis*, several of these open questions now become accessible.

## Experimental Procedures

### Strain construction

General methods for strain construction were performed according to published protocols (62, 63). Molecular cloning was performed using the Gibson assembly method with HiFi assembly enzyme mix (NEB) (64). PCR amplification templates were derived from either Chromosomal DNA isolated from the prototrophic domesticated *B. subtilis* strain PY79 or plasmid pKRH83. Introduction of DNA into PY79 derivatives was conducted by transformation (65). The bacterial strains, plasmids, and oligonucleotide primers used in this study are listed in Supplemental Tables 1-3 respectively.

### Media and growth conditions

For general propagation, *B. subtilis* strains were grown at 37°C in Lennox lysogeny broth (LB) medium (10 g tryptone per liter, 5 g yeast extract per liter, 5 g NaCl per liter) or on LB plates containing 1.5% Bacto agar. Where indicated, *B. subtilis* strains were grown in the nutrient-limiting medium Medium for Competence with 2% fructose (MC-Fru ; 61.5 mM K_2_HPO_4_, 38.2 mM KH_2_PO_4_, 2% (w/v ; 110 mM) D-fructose, 3 mM Na_3_C_6_H_5_O_7_ · 2H_2_0, 80 µM ferric ammonium citrate, 0.1% (w/v) casein hydrolysate, 11 mM L-Glutamic acid potassium salt monohydrate) substituted for 2% glucose to prevent catabolite repression of *P_xyl_*promoter (62). When appropriate, antibiotics were included in the growth medium as follows: 100 μg mL^-1^ spectinomycin, 5 μg mL^-1^ chloramphenicol, 5 μg mL^-1^ kanamycin, 10 μg mL^-1^ tetracycline, and 1 μg mL^-1^ erythromycin plus 25 μg mL^-1^ lincomycin (mls). When required, 0.5% (w/v ; ∼30 mM) D-xylose was added to cultures to induce protein expression.

### Producing lysed protoplasts for B. subtilis transformation

A single colony of the *B. subtilis* strain bBB050 (Cm^R^) was inoculated into LB medium and incubated at 37 °C with 250 rpm shaking for 3 hr. The OD_600_ of a 1:10 dilution of the culture was measured, and then 1 mL of culture was pelleted at 21,000 x G – 2 min and the supernatant removed completely. The pellet was resuspended to an OD_600_ = 10 in *Bacillus* protoplasting buffer (50 mM tris pH 8.0, 50 mM NaCl, 5 mM MgCl_2_, 25% (w/v) sucrose, 0.2 mg/mL lysozyme) and incubated in a 37 °C water bath for 30 min to protoplast cells. The sample was removed from the water bath and left at room temperature until transformation, at which time protoplasts were pelleted at 10,000 x G for 5 min, the supernatant was completely removed, and then an equal volume of ddH_2_O was added. The protoplasts were lysed by resuspension in the ddH_2_O by repeated pipetting.

### Transformation efficiency assays

Single colonies of *B. subtilis* strains of interest were inoculated into MC-Fru and cultured at 37 °C with 250 rpm shaking until an OD_600_ = 0.2 – 0.5 was reached. Cells were pelleted at 10,000 x G – 2 min and resuspended in ∼15% of residual supernatant to concentrate cells, OD_600_ of a 1:20 dilution of resuspended cells was measured, and the resuspensions were diluted into fresh MC-Fru to an OD_600_ = 0.05. Cultures were incubated at 37 °C with 250 rpm shaking for 2 hr (1 hr prior to max natural competence induction in these conditions), and strains containing *P_xyl_-comK* were induced with 0.5% xylose to maximize the proportion of competent cells in the populations. The strains continued to be cultured at 37 °C with 250 rpm shaking for 1 hr to allow for maximal natural competence induction, and then ∼ 10^5^ lysed Cm^R^ protoplasts per µL competent cells were added to the cultures (31). Transformation was allowed to proceed under the same culturing conditions for 2 hr. 10-fold serial dilutions of each sample were made down to 10^6^-fold diluted in PBS, and appropriate dilutions were plated onto LB and LB – Cm^5^ agar plates and incubated for 16-20 hr at 37 °C to allow for colony growth. The number of transformants and total cells were calculated from the single colonies on LB – Cm^5^ and LB respectively, and the transformation efficiency was calculated as the ratio of transformants to total cells in a given sample.

### ComGC Western blotting

*B. subtilis* strains of interest were cultured according to the same protocol noted for transformation efficiency assays, but 1 mL of culture was centrifuged at 21,000 x G for 2 min to pellet cells after 3 hr incubation post-dilution, and supernatant was completely removed. The OD_600_ of a 1:10 dilution of each culture was measured, and each cell pellet was resuspended in cell lysis buffer (25 mM Tris pH 8.0, 25 mM NaCl, 3 mM MgCl_2_, 1 mM CaCl_2_, 0.2 mg/mL lysozyme, 0.1 mg/mL DNase I) to an OD_600_ = 10 based on the previous measurements. Resuspended cells were incubated in a 37 °C water bath for 20 minutes to lyse cells and degrade genomic DNA. Cell lysates were mixed with an equal volume of 2X reducing tricine sample buffer (200 mM tris pH 6.8, 40% (v/v) glycerol, 2% (w/v) SDS, 0.04% (w/v) Coomassie Blue G-250, 2% (v/v) beta-mercaptoethanol) and heated to 37 °C for 30 minutes for protein denaturation. 5 µL of each preparation was added to wells of a tris-tricine mini gel (Stacking gel: 1M tris pH 8.45, 4% (w/v) acrylamide/bis-acrylamide (29:1) ; Resolving gel: 1M tris pH 8.45, 15% (w/v) glycerol, 10% (w/v) acrylamide/bis-acrylamide (29:1)) and electrophoresed (Running buffer: 0.1 M tris-Cl, 0.1 M tricine, 0.1% (w/v) SDS) at 100 V until loading dye exited the gel (typically 1.75 – 2 hr). Separated proteins were Western transferred to a 0.2 µm PVDF membrane using the semi-dry transfer method at 15V for 20 min with Towbin transfer buffer (66). The membrane was washed 3X in ddH_2_O for 5 min with gentle shaking to remove transfer buffer, then the membrane was stained with 0.1% Ponceau S solution to detect total protein for loading normalization. The membrane was blocked with 5% (w/v) milk in TBS-T for 1 hr with gentle shaking, and then the membrane was incubated with 1:2000 rabbit-derived antisera containing 1° ComGC antibody (TBS-T + 1% (w/v) milk) overnight at 4 °C (37). The membrane was washed 3X in TBS-T for 5 min with gentle shaking, and then the membrane was incubated with 1:20,000 Abcam goat anti-rabbit 2° HRP antibody (TBS-T + 1% (w/v) milk) for 1 hr at room temperature with gentle shaking. The membrane was washed as described previously, and then developed using Clarity ECL substrate (Bio-Rad) and imaged for chemiluminescence using a Bio-Rad ChemiDoc Touch imaging system.

### Preparing cover glass and agar pads for use in microscopy

All cover glass used in microscopy experiments was pre-cleaned prior to use. 22 mm x 22 mm #1.5 borosilicate coverslips (DOT Scientific) were placed in a Wash-N-Dry™ coverslip rack (Sigma) and submerged in ∼80 mL of 1 M NaOH in a 100 mL glass beaker. The beaker was then placed into an ultrasonic cleaning bath (frequency = 40 kHz, power = 120W) and sonicated for 30 min to remove the thin grease layer present on the coverslips. The coverslip rack was submerged into a fresh 250 mL glass beaker filled with ddH_2_O, the ddH_2_O was removed, and then the coverslips were washed 3X with 250 mL of ddH_2_O. The coverslips were then either air-dried overnight or dried immediately with compressed air.

For the preparation of agar pads, pre-cleaned borosilicate glass microscope slides were first rinsed free of detritus using ddH_2_O and were then either air-dried overnight or dried immediately with compressed air. Working in a fume hood, half of the total rinsed slides were dipped into a 2% (v/v) solution of dichlorodimethylsilane (in chloroform) so that their entire surface was contacted by the solution. The solution was allowed to drip off the slides into the original container, and the remaining organic solvent on the slides was allowed to evaporate in the fume hood for 30 min. The slides were then thoroughly rinsed in ddH_2_O and dried as mentioned previously to generate dry glass slides with extremely hydrophobic surfaces. Two pieces of lab tape (VWR #89098-074) were placed on top of one another, running lengthwise across an unrinsed microscope slide. For every agar pad to be produced, 2 of these taped slides were made. ∼15 minutes prior to imaging, conditioned MC-Fru medium (0.22 µm PES filter sterilized) from cultures grown to competence via the transformation efficiency assay protocol was heated to 37 °C, added 1:1 to 90 °C 2% (w/v) molten LE agarose (SeaKem) in ddH_2_O, and then vortexed to make molten 1% (w/v) agarose in 0.5X conditioned MC-Fru medium. 20 µL of this mixture was applied to the surface of a hydrophobic glass slide, and a solidified pad was produced according to a previously established protocol, using a rinsed (but untreated and hydrophilic) glass slide to form the top of the pad (67). This specific setup allows for the agar pad to stick to the bottom slide and easily slide out from the top slide, leaving an unmarred and flat surface for imaging.

### Fluorescent labeling – ComGC pili

For all experiments involving ComGC pilus labeling, *B. subtilis* strains of interest were cultured according to the same protocol noted for transformation efficiency assays. After 3 hr incubation post-dilution, 100 uL of each culture was transferred to a 37 °C pre-warmed 13 mm glass test tube, and 25 ug/mL Alexa Fluor 488-maleimide was added to each culture aliquot. The aliquots were incuabted in the dark at 37 °C on a rolling drum for 20 minutes to allow for ComGC^Cys^ pilin labeling. The aliquots were transferred to centrifuge tubes, centrifuged at 5,000 x G for 30 seconds to gently pellet cells, and all supernatant was removed. The cell pellets were washed by gently resuspending in conditioned MC-Fru medium (0.22 µm PES filter sterilized) via gentle pipetting. The cells were centrifuged again at 5,000 x G for 30 seconds, the supernatant removed, and the cells gently resuspended in one-tenth the original volume of conditioned MC-Fru medium.

### Fluorescent labeling – PCR product

An ∼4.5 kb PCR product was amplified from genomic DNA of a *B. subtilis* strain bearing *P_hysp_-bdbDC* at the *ycgO* locus (bJZ185) using LongAmp Taq DNA Polymerase (NEB) and oligonucleotide primer pair (oJZ436 + oJZ437). This PCR product was purified using the E.Z.N.A. Cycle Pure Kit (Omega Bio-Tek) following the manufacturer’s instructions. 1 µg of PCR product was fluorescently labeled using the *Label* IT-Cy5 Nucleic Acid Labeling Kit (Mirus) following the manufacturer’s instructions, with a sufficient quantity of *Label* IT-Cy5 reagent to covalently link a fluorophore to between 1 in 20 and 1 in 60 bases (75 – 230 Cy5 molecules per DNA molecule). The Cy5 labeled PCR product was purified using the E.Z.N.A. Cycle Pure Kit as mentioned previously. The final product was electrophoresed on a 1% agarose in TAE mini gel at 100 V for 45 min, stained using SYBR Safe dye (Invitrogen) according to the manufacturer’s instructions, and quantified by densitometric analysis of the product band compared to a reference band of known quantity.

### Microscopy – Alexa Fluor 488-maleimide labeled ComGC pili and Label IT-Cy5 labeled PCR product

For the initial identification of ComGC pili, quantification of pilus production, pilus length measurements, and the determination of ComGC pilus retraction rates, cells were labeled with Alexa Fluor 488-maleimide as previously described. 0.5 µL of resuspended, labeled cells was applied to the center of a conditioned MC-Fru agar pad, and a pre-cleaned 22 mm x 22 mm #1.5 borosilicate coverslip was applied to the drop of culture. The coverslip was gently compressed with a gloved finger to ensure cells made contact with the agar pad, and then the space between the coverslip and glass slide was sealed by applying molten (60 °C) VaLAP sealant (1:1:1 petroleum jelly:lanolin:paraffin) to the coverslip edges. The cells were imaged using a Zeiss Axio Observer.Z1 inverted epifluorescence microscope equipped with a Zeiss Plan-Apochromat 100X/1.4 Oil PH3 objective lens and and a Teledyne Photometrics CoolSNAP HQ^2^ CCD camera. Cells were typically exposed to light from a Zeiss Colibri 469 nm LED module with Zeiss filter set 38 HE for 2 s at 100% LED power to observe ComGC pili. Cell bodies were imaged using phase-contrast microscopy (50 ms exposures). Imaging was performed at 37 °C, achieved by a PECON Heater S objective heater calibrated via thermocouple. Time lapse microscopy was performed with the above conditions, with exposures occuring every 5 – 11 s depending on the experiment.

For co-localization experiments on ComGC pili and PCR products, cells were labeled with Alexa Fluor 488-maleimide as previously described. Just prior to deposition of labeled cells onto a conditioned MC-Fru agar pad, Cy5 labeled PCR product was added to the cell suspension to a final concentration of 120 pg/uL. The new mixture of Alexa Fluor 488 labeled cells and Cy5 labeled PCR product was applied to a conditioned MC-Fru agar pad as described above. imaging was performed using a Zeiss Axio Imager.Z2 upright epifluorescence microscope equipped with a Zeiss Plan-Apochromat 100X/1.4 Oil PH3 objective lens and a Teledyne Photometrics Prime 95B sCMOS camera. Cells were typically exposed to light from a Zeiss Colibri 469 nm LED module with Zeiss filter set 38 HE for 200 ms at 20% LED power to observe ComGC pili, while *Label* IT-Cy5 labeled PCR product was observed using light from a Zeiss Colibri 631 nm LED module with Zeiss filter set 90 HE for 500 ms at 20% LED power. Cell bodies were imaged using phase-contrast microscopy (100 ms exposures). Z-stacks were acquired in 0.2 µm increments starting 1 µm above the midcell plane and progressing until 1 µm below the midcell plane. Due to a fortuitous imaging delay during phase-contrast acquisition, the time interval between epifluorescent image acquisition was ∼ 5 s, producing a time lapse.

### Microscopy – localization of GFP-ComEA and GFP-ComFA

Fluorescence microscopy was performed as previously described(68, 69). Exposure times were typically 500 ms for GFP-ComEA and 250 ms for GFP-ComFA. Membranes of *gfp-comEA* cells were stained with TMA-DPH (Molecular Probes), at a final concentration of 0.01 mM, and imaged with exposure times of 200 ms. Cell bodies of *gfp-comFA* cells were imaged using phase-contrast microscopy (20 ms exposures). Fluorescence images were analyzed, adjusted, and cropped using Metamorph v 6.1 software (Molecular Devices).

### Microscopy – post-processing of Alexa Fluor 488-maleimide labeled ComGC pili and Label IT-Cy5 labeled PCR product images

All epifluorescence microscopy images presented in Fig. 2 through Fig. 4 were deconvoluted prior to publication. First, point-spread-functions (PSFs) were computationally estimated for Alexa Fluor 488 and Cy5 signals using the PSF Generator plugin (EPFL Biomedical Imaging Group) on Fiji (NIH) (70). For Alexa Fluor 488 signal captured on the Zeiss Axio Observer.Z1 microscope configured as described above, the following parameters were entered into the PSF Generator: Optical model = Born and Wolf 3D Optical Model ; Refractive index immersion = 1.518 (corresponding to Immersol 518F) ; Accuracy computation = Best ; Wavelength = 516 nm (corresponding to Alexa Fluor 488 emission maximum) ; NA (numerical aperture) = 1.4 ; Pixelsize XY = 65 nm ; Z-step = 250 nm ; Size XYZ – X = 256, Y = 256, Z = 65 ; Display = Linear, 16-bits, Fire. The Z-stack composing the PSF was Z-projected to a single 256 x 256 pixel image by averaging the pixel intensities of each individual pixel in the 256 x 256 pixel array across the 65 Z-slices of the Z-stack, resulting in a 2D PSF. For Alexa Fluor 488 signal captured on the Zeiss Axio Imager.Z2 microscope configured as described above, the same procedure was performed to estimate the PSF, only changing Pixelsize XY to 110 nm. For Cy5 signal captured on the Zeiss Axio Imager.Z2 microscope configured as described above, all parameters were kept constant to estimate the PSF, only changing Wavelength = 666 nm (corresponding to Cy5 emission maximum).

Epifluorescence microscopy images were deconvolved using the DeconvolutionLab2 plugin (EPFL Biomedical Imaging Group) for Fiji (NIH). Individual epifluorescence microscopy images were opened in Fiji, and the DeconvolutionLab2 plugin was run. The PSF generated as above, corresponding to a particular epifluorescence signal, was used to deconvolve that signal with DeconvolutionLab2 (Algorithm = Richardson-Lucy, 100 iterations).

### Quantification of pili produced in comGC merodiploid strains

Images of Alexa Fluor 488-maleimide labeled cells from each merodiploid strain, acquired as described previously, were opened in the Fiji image processing package (ImageJ2, NIH). The plugin ObjectJ was first used to identify 300 individual cells of each strain from phase-contrast images. Narrow filaments of length greater than 0.5 µm that were directly connected to the cell bodies of these 300 cells, which were surmised to be ComGC pili, were identified and counted in the green epifluorescence channel. The extracellular space immediately adjacent to the 300 cell bodies was scanned in the green epifluorescence channel for foci, which were likely sheared ComGC pili, and foci within 0.5 µm of the cell body were counted. These epifluorescence data were graphed using Microsoft Excel.

### Measurement of ComGC pili lengths

Images of Alexa Fluor 488-maleimide labeled cells of the *comGC^E56C^*and *comGC^S65C^* merodiploid strains, acquired as described previously, were opened in Fiji (NIH). The images were scaled up 5-fold using bilinear interpolation, and the phase-contrast and green epifluorescence image channels were split for each pilus-producing cell. Each channel was converted into a binary image using Fiji’s default thresholding parameters, and then outlines of the signal present in each channel were produced. The outline view of the phase-contrast channel demarcated the cell boundary and extracellular space, while the green epifluorescence channel outline divided a pilus from the extracellular space. These outlines were merged together to simultaneously display both the cell boundary and pilus boundary. Pilus length was manually measured by placing the start of a segmented line on the cell boundary, approximately where the medial axis of the pilus would cross the cell boundary, and creating a line that roughly followed the pilus medial axis to the extreme tip of the pilus.

### ComGC pilus retraction rate measurements

Time lapse microscopy images of Alexa Fluor 488-maleimide labeled *comGC^S65C^* cells, acquired as described previously, were visually scanned for potential ComGC pilus retraction events using Fiji (NIH). Once identified, cells with retracting pili were isolated, and pilus length was measured for each time point of the time lapse as described previously. Frame-to-frame retraction rates were calculated by dividing the change in pilus length between frames by the time interval of the time lapse. These data were collected for ten individual pilus retraction events, which included 44 instances of frame-to-frame retraction. Microsoft Excel was used to perform the t-test and ANOVA (using the Real Statistics Resource Pack release 8.6.3) referenced in the results section.

### Localization analysis of fluorescent proteins (ComGC, ComEA, ComFA) and PCR product

Images acquired as previously described for Alexa Fluor 488-maleimide labeled *comGC^S65C^* merodiploid, *gfp-comEA*, and *gfp-comFA* strains were opened in Fiji (NIH). For labeled ComGC^S65C^ pili, individual pilus-producing cells were isolated and identified in the phase-contrast channel, and a 4x4 pixel white square was added to the green epifluorescence channel where the pilus medial axis would cross the cell boundary to mark the base of pilus production. Because maleimide labeling also resulted in cell membrane staining, cell bodies were checked against the green fluorescence channel to identify cell septa not visible in phase-contrast images. If a septum was identified, a 1-pixel wide white line was added onto the phase-contrast image across the cell body to allow MicrobeJ to detect multiple cells in the image; this process was repeated for 232 individual pilus production events. MicrobeJ (version 5.11x) was used to define cell bodies and medial axes from phase-contrast images, and pili bases were identified from the green epifluorescence images by adjusting the plugin’s sensitivity parameters (tolerance and Z-score) until only the added white square was detected as a fluorescent focus (71). The identified cells and pili boundaries were associated with one another, and a heatmap of pilus production localization on a size-normalized rod-shaped cell was made within the “Heatmap(s)” tab of the results window. This heatmap was scaled up 10-fold using bilinear interpolation to create the final version included in Fig. 5.

For localization of GFP-ComEA foci, cell bodies were identified by TMA-DPH epifluorescence signal, and the outline of each cell was filled in using a white rounded rectangle with a 6-pixel wide black border to enhance MicrobeJ detection of cells. GFP-ComEA foci were identified in the green epifluorescence channel and marked with a 4x4 pixel white square to enhance contrast. Microbe J (version 5.11x) was used as described above to generate a heatmap of GFP-ComEA localization. For localization of GFP-ComFA foci, cell bodies of isolated single cells were identified using phase-contrast images, and GFP-ComFA foci were identified in the green epifluorescence channel. A localization heatmap was generated as described previously. For localization of Cy5-labeled PCR product binding to Alexa Fluor 488-maleimide labeled cells, time course images acquired as described previously were opened in Fiji (NIH). a binding event was defined as the co-localization of a Cy5 focus with either the cell body or a labeled ComGC filament for at least 3 consecutive frames (≥ 15 s of co-localization). Cell bodies were identified as described previously, and Cy5-labeled PCR product foci were identified in the red epifluorescence channel and contrast-enhanced with a 4x4 white square as described previously. A localization heatmap was then generated as described previously.

## Acknowledgments

The authors thank Daniel B. Kearns for supplying us with pKRH83 (*lacA::P_comG_-comGC*) used in the mutagenesis and construction of *comGC^Cys^* merodiploid strains, Thorsten Mascher for providing us with the sequences for pBS0E and pBS0E-*P_xyl_*which was used in the construction of the *P_xyl_-rbs-comK* allele for natural competence induction, Jonathan Lombardino for constructing the violin plots found in Fig. 2C, and the entire Burton laboratory for thoughtful criticism, discussion, and comments. This work was supported by the Rita Allen Foundation Milton E. Cassel Award. JZ was supported by an NIH T32 training grant (GM07215).

## Notes

### Competing Interest Statement

The authors have declared no competing interest.

