## Supplemental Information for "Visualizing dynamic competence pili and DNA capture throughout the long axis of *Bacillus subtilis*"


Running Title: Distribution of *B. subtilis* pili and DNA capture


Keywords: competence, natural transformation, pili, *B. subtilis*



* Correspondence:
, 608-890-0510

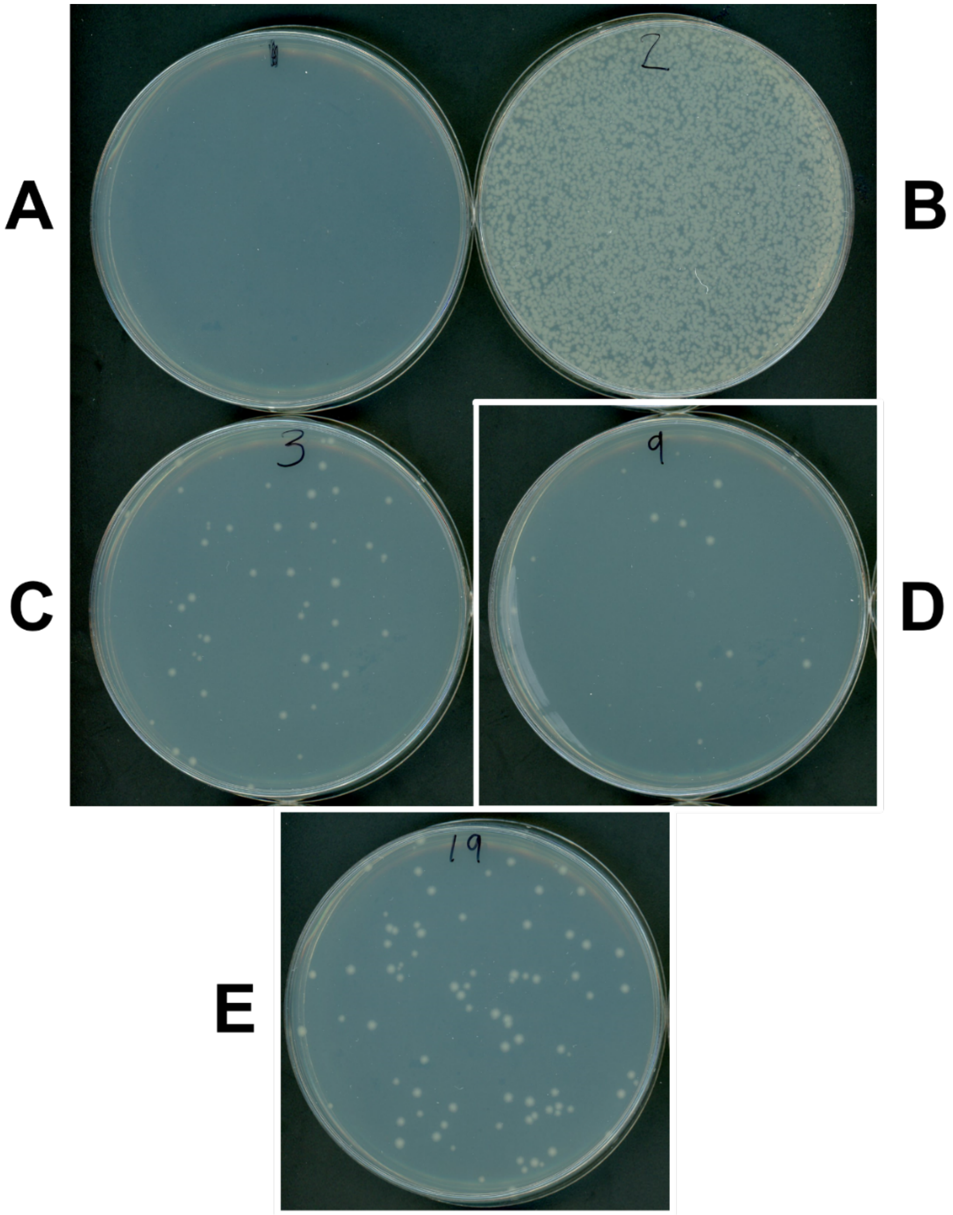


**Supplemental Figure 1.**
Select $ComGC^{Cys}$ variants support low -efficiency transformation as the sole ComGC variant produced. Transformants resulting from transformation of *B. subtilis* genomic DNA bearing a spectinomycin resistance gene and plating on LB + Spc, using strains (A) PY79 (no DNA) (B) PY79 (C) *lacA::*$P_{comG}$*-comGC*$^{E56C}$ ; Δ*comGC*$^{WT}$ (D) *lacA::*$P_{comG}$*-comGC*$^{S65C}$ ; Δ*comGC*$^{WT}$ (E) *lacA::*$P_{comG}$*-comGC*$^{WT}$ ; Δ*comGC*$^{WT}$.

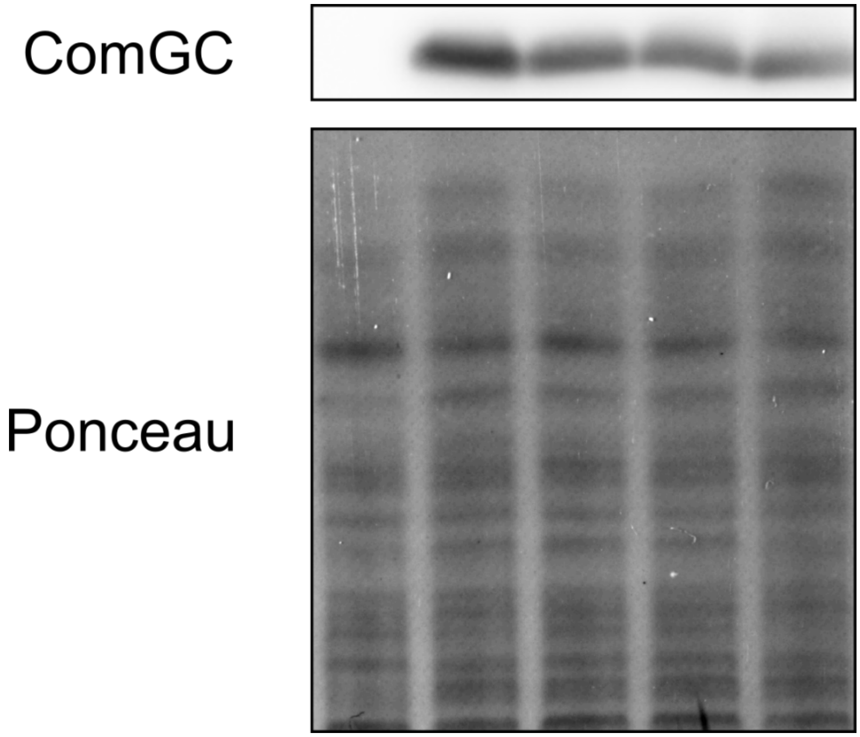


**Supplemental Figure 2.**
Representative whole cell lysate ComGC Western blots and Ponceau total protein staining of the following strains cultured until maximum competence was induced, from left to right: *ΔcomK* (uninduced), *comGC*$^{WT}$, *comGC*$^{E56C}$, *comGC*$^{S65C}$, *ΔcomK*. All strains ectopically maintained *P*$_{xyl}$*-rbs-comK* and were induced with 0.5% (w/v) xylose unless otherwise noted.

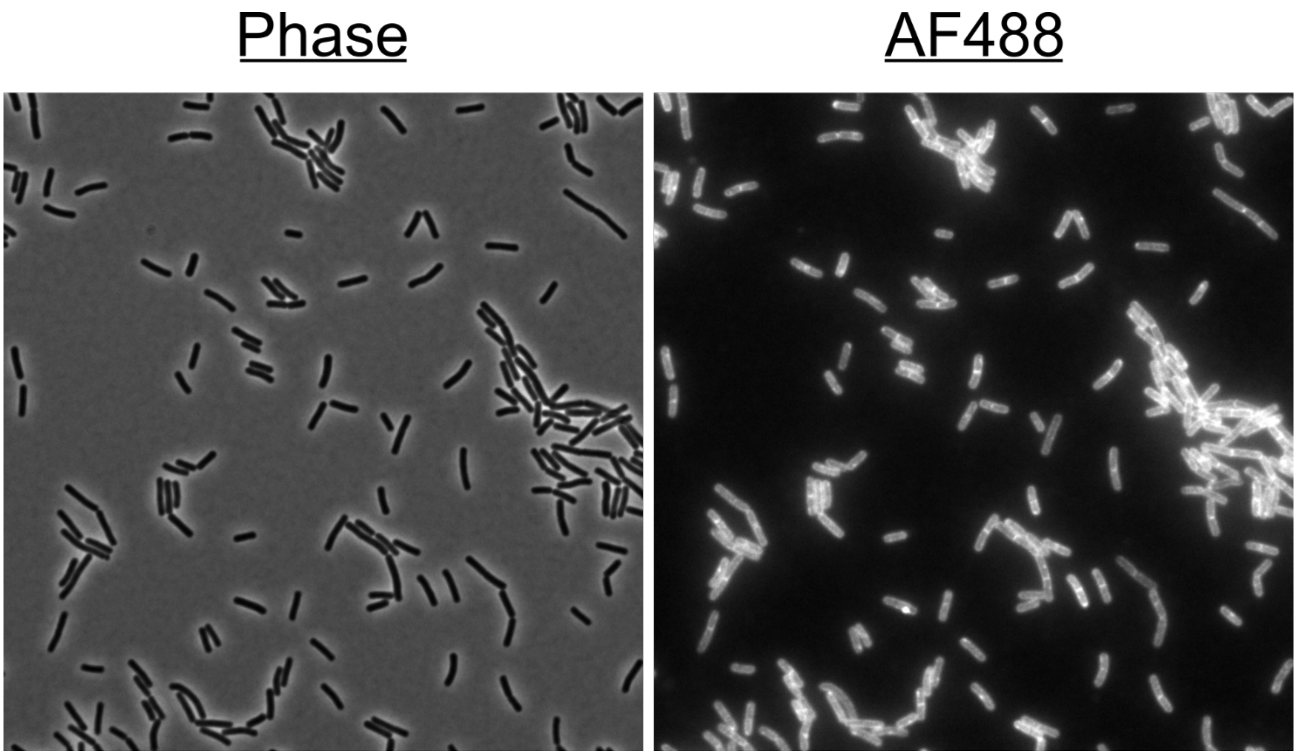


**Supplemental Figure 3.**
Representative imaging field of maleimide-labeled *comGC*$^{WT}$. No external protruding structures were observed.

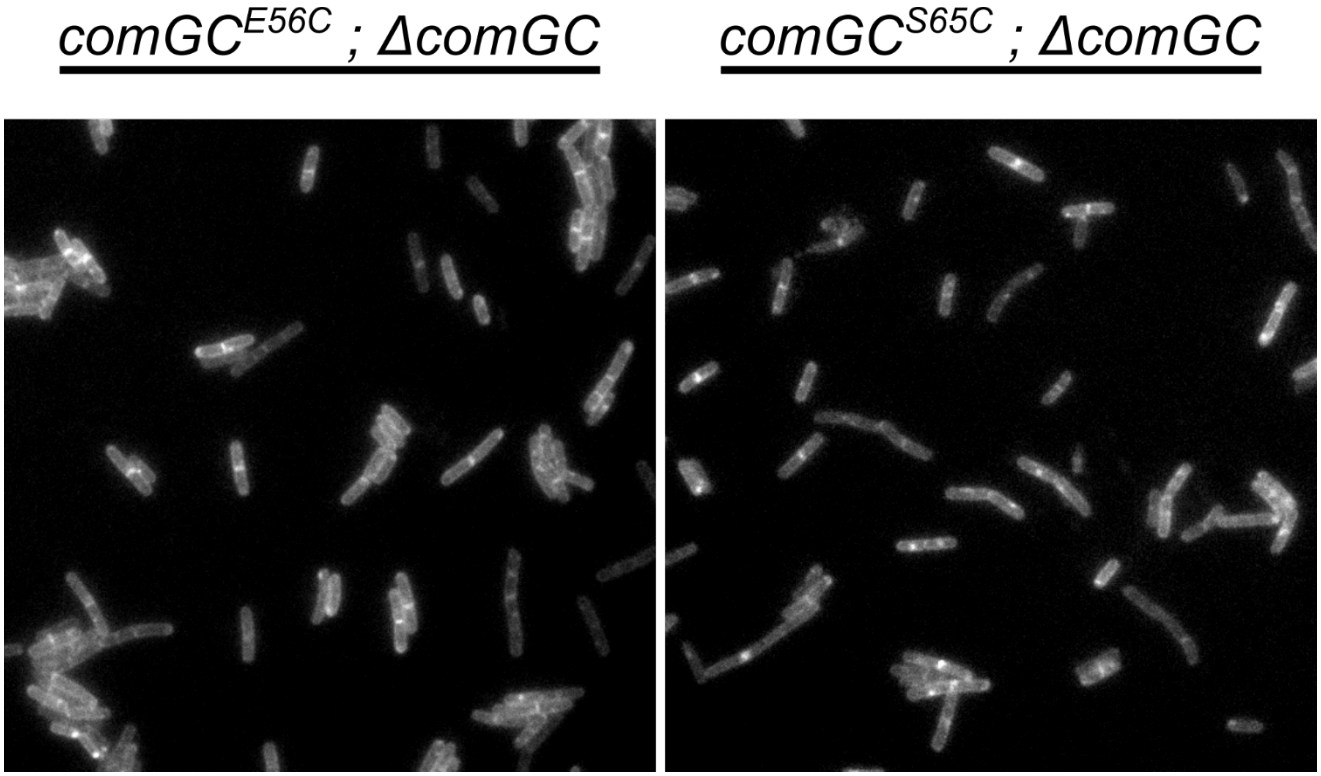


**Supplemental Figure 4.**
Representative imaging field of maleimide-labeled cells with either *comGC*$^{E56C}$ or *comGC*$^{S65C}$ as the sole copy of *comGC* expressed. No external protruding structures were observed in the AF488 channel.

**Supplemental Movie 1.**

Example of a ComGC$^{S65C}$ pilus binding to fluorescently labeled DNA

**Supplemental Movie 2.**

Example of a ComG$C^{S65C}$ pilus retracting

**Supplemental Table 1.** Strains used in this study

| Strain | Genotype | Parent strain | Source, Reference |
|---|---|---|---|
| DH5α | F⁻ *endA1 glnV44 thi1 recA1 relA1 gyrA96 deoR nupG purB20* ϕ80d *lacZ*ΔM15 Δ(*lacZYA-argF*)U169, hsdR17($r_K^- m_K^+$), λ⁻ | | (1) |
| eJZ040 | pJZ026 (*yhdGH::*$P_{xyl(-CRE)}$ *(kan)*) | DH5α | This work |
| eJZ064 | pJZ049 (*yhdGH::*$P_{xyl(-CRE)}$*-rbs-comGC (kan)*) | DH5α | This work |
| eJZ095 | pJZ069 (*yhdGH::*$P_{xyl}$*-rbs-comK (kan)*) | DH5α | This work |
| PY79 | Prototrophic domesticated laboratory strain | | (2) |
| bBB036 | *comK::spc* | PY79 | This work |
| bBB050 | *amyE::*$P_{comE}$*-gfp-comEA (cat)* | PY79 | This work |
| bSC050 | *yvbJ::*$P_{comF}$*-gfp-comFA (erm)* | PY79 | (3) |
| bJZ083 | *comGC::kan* | 168 | (3) |
| bJZ096 | *lacA::*$P_{comG}$*-comGC*$^{E56C}$ *(erm)* | PY79 | This work |
| bJZ103 | *lacA::*$P_{comG}$*-comGC*$^{S65C}$ *(erm)* | PY79 | This work |
| bJZ117 | *lacA::*$P_{comG}$*-comGC*$^{WT}$ *(erm)* | PY79 | This work |
| bJZ140 | *lacA::*$P_{comG}$*-comGC*$^{E56C}$ *(erm) ; comGC::kan* | bJZ096 | This work |
| bJZ146 | *lacA::*$P_{comG}$*-comGC*$^{S65C}$ *(erm) ; comGC::kan* | bJZ103 | This work |
| bJZ156 | *lacA::*$P_{comG}$*-comGC*$^{WT}$ *(erm) ; comGC::kan* | bJZ117 | This work |
| bJZ405 | *yhdGH::*$P_{xyl}$*-rbs-comK (kan)* | PY79 | This work |
| bJZ407 | *yhdGH::*$P_{xyl}$*-rbs-comK (kan) ; comK::spc* | bJZ405 | This work |
| bJZ418 | *yhdGH::*$P_{xyl}$*-rbs-comK (kan) ; amyE::*$P_{comE}$*-gfp-comEA (cat)* | bJZ405 | This work |
| bJZ419 | *yhdGH::*$P_{xyl}$*-rbs-comK (kan) ; lacA::*$P_{comG}$*-comGC*$^{E56C}$ *(erm)* | bJZ405 | This work |
| bJZ423 | *yhdGH::*$P_{xyl}$*-rbs-comK (kan) ; lacA::*$P_{comG}$*-comGC*$^{S65C}$ *(erm)* | bJZ405 | This work |
| bJZ427 | *yhdGH::*$P_{xyl}$*-rbs-comK (kan) ; lacA::*$P_{comG}$*-comGC*$^{WT}$ *(erm)* | bJZ405 | This work |

**Strain construction:**

Note: All strains below are isogenic with PY79, unless otherwise listed.

**eJZ040** [pJZ026 (*yhdGH::*$P_{xyl(-CRE)}$ *(kan)*)] was generated by transformation of Gibson assembled **pJZ026** into chemically competent **DH5α** via heat-shock, selecting for $Amp^R$ (4). Transformants were verified by colony PCR screening.

**eJZ064** [pJZ049 (*yhdGH::*$P_{xyl(-CRE)}$*-rbs-comGC (kan)*)] was generated by transformation of Gibson assembled **pJZ049** into chemically competent **DH5α** via heat-shock, selecting for $Amp^R$ (4). Transformants were verified by colony PCR screening.

**eJZ095** [pJZ069 (*yhdGH::*$P_{xyl}$*-rbs-comK (kan))*] was generated by transformation of Gibson assembled **pJZ069** into chemically competent **DH5α** via heat-shock, selecting for $Amp^R$ (4). Transformants were verified by colony PCR screening.

**bBB036** [*comK::spc*] was generated by transformation of BD2259 (5) genomic DNA into competent **PY79** selecting $Spc^R$, followed by two successive backcrosses of transformant genomic DNA into competent **PY79** selecting $Spc^R$.

**bBB050** [*amyE::PcomE-gfp-comEA (cat)*] was generated by a double crossover of **pBB401** into competent **PY79** selecting $Cm^R$. The double crossover was confirmed by PCR from genomic DNA.

**bJZ096** [*lacA::*$P_{comG}$*-comGC*$^{E56C}$ *(erm)*] was generated first by overlap extension PCR (6) that produced *lacA::*$P_{comG}$*-comGC*$^{E56C}$ *(erm)*. Primers pairs (**oJZ149** + **oJZ152**) and (**oJZ150** + **oJZ151**) were used in conventional PCR with **pKRH83** to generate two *lacA::*$P_{comG}$*-comGC*$^{E56C}$ *(erm)* fragments with two homologous ends. The products of each reaction were isolated and extracted from an agarose gel (4), and an equimolar ratio of each purified product was used in a subsequent PCR (without primers) to make full length *lacA::*$P_{comG}$*-comGC*$^{E56C}$ *(erm)*. Primer pair (**oJZ149** +

**oJZ150**) was added approximately halfway through the reaction to begin exponential amplification of full length *lacA::P$_{comG}$-comGC$^{E56C}$ (erm)*. Once amplification was verified by agarose gel electrophoresis, full length *lacA::P$_{comG}$-comGC$^{E56C}$ (erm)* PCR product was used to directly transform competent **PY79** selecting MLS$^R$. Double crossover recombination was verified by PCR of genomic DNA using primer pair (**oJZ212** + **oJZ213**) and the same PCR product was Sanger sequenced (primer **oJZ214**) to verify that no mutations other than *comGC$^{E56C}$* had been introduced.

**bJZ103** [*lacA::P$_{comG}$-comGC$^{S65C}$ (erm)*] was generated as described for **bJZ096**, but the conventional PCR primer pairs were swapped to (**oJZ149** + **oJZ176**) and (**oJZ150** + **oJZ175**).

**bJZ117** [*lacA::P$_{comG}$-comGC$^{WT}$ (erm)*] was generated as described for **bJZ096**, but the conventional PCR primer pairs were swapped to (**oJZ149** + **oJZ202**) and (**oJZ150** + **oJZ201**).

**bJZ140** [*lacA::P$_{comG}$-comGC$^{E56C}$ (erm) ; comGC::kan*] was generated by transformation of competent **bJZ096** with *comGC::kan* PCR product (primer pair **oJZ223** + **oJZ225** with **bJZ083** genomic DNA template) selecting Kan$^R$ followed by screening 10 transformants for MLS$^R$ by patching single colonies. Double crossover recombination of comGC::kan was confirmed by PCR using primer pair (**oJZ199** + **oJZ200**) and strain genomic DNA.

**bJZ146** [*lacA::P$_{comG}$-comGC$^{S65C}$ (erm) ; comGC::kan*] was generated as described for **bJZ140**, but competent **bJZ103** was transformed with *comGC::kan* PCR product

**bJZ156** [*lacA::P$_{comG}$-comGC$^{WT}$ (erm) ; comGC::kan*] was generated as described for **bJZ140**, but competent **bJZ117** was transformed with *comGC::kan* PCR product

**bJZ405** [*yhdGH::P$_{xyl}$-rbs-comK (kan)*] was generated first by preparing **pJZ069** plasmid DNA from **eJZ095** by alkaline lysis (4). **pJZ069** was then restriction digested with ScaI-HF to linearize (to facilitate double crossover recombination), and then transformed into competent **PY79** selecting for Kan$^R$. The transformed allele was tested for functionality by complementation in **bJZ407**.

**bJZ407** [*yhdGH::P$_{xyl}$-rbs-comK (kan) ; comK::spc*] was generated by transforming competent **bJZ405** with genomic DNA from **bBB036** and selecting for Spc$^R$ followed by screening 10 transformants for Kan$^R$ by patching single colonies. Kan$^R$ Spc$^R$ transformants were tested for *comK* complementation by transformation of induced (0.5% (w/v) xylose) and non-induced cultures with lysed **bBB050** protoplasts (genomic DNA carrying *amyE::PcomE-gfp-comEA (cat)*) and selecting for Cm$^R$. All transformants tested were non-transformable when not induced, and fully transformable when induced, indicating full complementation of *comK::spc*.

**bJZ418** [*yhdGH::P$_{xyl}$-rbs-comK (kan) ; amyE::P$_{comE}$-gfp-comEA (cat)*] was generated by transforming competent **bJZ405** with genomic DNA from **bBB050** selecting for Cm$^R$. Transformants were expected to express *gfp-comEA* in a much greater proportion of cells when induced with 0.5% xylose, and this was confirmed by epifluorescence microscopy.

**bJZ419** [*yhdGH::P$_{xyl}$-rbs-comK (kan) ; lacA::P$_{comG}$-comGC$^{E56C}$ (erm)*] was generated by transforming competent **bJZ405** with genomic DNA from **bJZ096** selecting for MLS$^R$. Double crossover recombination was verified by PCR with primer pair (oBOSE397 + oBOSE400) with strain genomic DNA as template.

**bJZ423** [*yhdGH::P$_{xyl}$-rbs-comK (kan) ; lacA::P$_{comG}$-comGC$^{S65C}$ (erm)*] was generated as described for **bJZ419**, but genomic DNA from **bJZ103** was used in the transformation.

**bJZ427** [*yhdGH::P$_{xyl}$-rbs-comK (kan) ; lacA::P$_{comG}$-comGC$^{WT}$ (erm)*] was generated as described for **bJZ419**, but genomic DNA from **bJZ117** was used in the transformation.

**Supplemental Table 2.** Plasmids used in this study

| Plasmid | Description | Source |
|---|---|---|
| pBB401 | *amyE::PcomE-gfp-comEA(cat)* | This work |
| pBS0E-XylR-$P_{xylA\ (V2)}$ | Empty replicative *B. subtilis* plasmid carrying *xylR*-$P_{xylA\ (V2)}$ | (7) |
| pKL147 | *dnaX*$^{3'}$*-gfp* | (8) |
| pKRH83 | *lacA::*$P_{comG}$*-comGC* | D. Kearns |
| pNS008 | *amyE::*$P_{SpoIIQ}$*-yfp (cat)* | D. Rudner |
| pJZ026 | *yhdGH::*$P_{xyl(-CRE)}$ *(kan)* | This work |
| pJZ049 | *yhdGH::*$P_{xyl(-CRE)}$*-rbs-comGC (kan)* | This work |
| pJZ069 | *yhdGH::*$P_{xyl}$*-rbs-comK (kan)* | This work |

**Plasmid Construction:**

**pBB401** [*amyE::PcomE-gfp-comEA(cat)*] was generated in a four-way ligation with an *Eco*RI-*Hind*III PCR product containing the promoter region for the *comE* operon (oligonucleotide primers oBMB116 and oBMB117 and genomic DNA from **PY79** as template), an *Hind*III-*Xho*I PCR product containing an optimized RBS and *gfp* (oligonucleotides oBMB127 and oBMB128 and **pKL147** as template), an *Xho*I-*Bam*HI PCR product containing *comEA*, all cloned into **pNS008** cut with *Eco*RI and *Bam*HI. **pNS008** was a gift from D. Rudner.

**pJZ026** [*yhdGH::*$P_{xyl(-CRE)}$ *(kan)*] was generated by a 2-fragment Gibson assembly. Primer pair 1 (**oJZ167** + **oJZ168**) and primer pair 2 (**oJZ169** + **oJZ170**) were used in separate PCRs with **pBOSE1408** as template in each reaction to amplify two fragments of **pBOSE1408** comprising almost the entire **pBOSE1408** sequence but lacking a catabolite repressive element found in the original $P_{xyl}$ cassette (9). Each PCR product was isolated and extracted from an agarose gel, and an equimolar ratio of each fragment was used to construct **pJZ026** by a 2-fragment Gibson assembly using HiFi master mix (NEB) according to the manufacturer's instructions (60 min incubation time).

**pJZ049** [*yhdGH::*$P_{xyl(-CRE)}$*-rbs-comGC (kan)*] was generated by a 2-fragment Gibson assembly. **pJZ026** was restriction digested with XhoI (NEB) to linearize for Gibson assembly. Primer pair 1 (**oJZ257** + **oJZ258**) was used in PCR with **pKRH83** as template to amplify *rbs-comGC* (**pKRH83** was a gift from D. Kearns). The amplified *rbs-comGC* product was extracted from an agarose gel and used as template in a subsequent PCR with primer pair 2 (**oJZ258** +**oJZ259**) to amplify *rbs-comGC* with appropriate homologous ends for Gibson assembly. XhoI-digested **pJZ026** and the subsequent *rbs-comGC* amplified product were extracted from an agarose gel, and an equimolar ratio of each fragment was used to construct **pJZ049** by a 2-fragment Gibson assembly using HiFi master mix (NEB) according to the manufacturer's instructions (60 min incubation time).

**pJZ069** [*yhdGH::*$P_{xyl}$*-rbs-comK (kan)*] was generated by a 3-fragment Gibson assembly. Primer pair 1 (**oJZ394** + **oJZ395**) was used in PCR with **pJZ049** to amplify the vector backbone and *yhdGH::kan* ectopic integration sequence, primer pair 2 (**oJZ396** + **oJZ397**) was used in PCR with **pBS0E-XylR-**$P_{xylA\ (V2)}$ (7) to amplify the $P_{xyl}$ *(xylR)* cassette, and primer pair 3 (**oJZ398** + **oJZ399**) was used in PCR with **PY79** genomic DNA to amplify *comK* including its transcriptional terminator. Each PCR product was isolated and extracted from an agarose gel, and an equimolar ratio of each fragment was used to construct **pJZ069** by a 3-fragment Gibson assembly using HiFi master mix (NEB) according to the manufacturer's instructions (60 min incubation time). The entire plasmid was then sequenced by Plasmidsaurus Oxford Nanopore sequencing for confirmation.

**Supplemental Table 3. Oligos used in this study**

| Primer | Sequence* |
|---|---|
| oBMB116 | GGAATTCCGCTTACGATGTCTGATGCAACC |
| oBMB117 | CGTAAGCTTCGCTTTCGTCATATTTTATTTTC |
| oBMB127 | GGCAAGCTTACATAAGGAGGAACTACTATGAGTAAAGGAGAAGAAC |
| oBMB128 | CGGCTCGAGTTTGTATAGTTCATCCATGC |
| oBOSE397 | CCAGCCAATATTCACTTGCTGCGG |
| oBOSE400 | CTCCCTGACCGCTCGGCTTCC |
| oJZ149 | GCTGCGCCTTATCCGGTAACTATCG |
| oJZ150 | CCAGCAACGCGGCCTTTTTACG |
| oJZ151 | *ATGACTGCATTTTGTCTTGATCATGA*AGGAC |
| oJZ152 | *TCATGATCAAGACAAAATGCAGTCAT*TTGTGC |
| oJZ167 | *GCATGCATTAGCTAGCATTA*CTCGAGACACTAGTATAAGC |
| oJZ168 | *ATTAAGCGCGGCGGGTGT*G |
| oJZ169 | *TAATGCTAGCTAATGCATGC*TTAGTTTGTTGCCCAAAC |
| oJZ170 | *ACACCCGCCGCGCTTAAT*G |
| oJZ175 | *GCCTTGCCTGTTTACAGT*CAGAGGGCTATG |
| oJZ176 | *ACTGTAAACAGGCAAGGC*TCGGAGTTTG |
| oJZ199 | TGGGCAACAAACTAAGCATGGACAATATGCGTTGAAAGGAGA |
| oJZ200 | CGAGTAATGCTAGCTAATGCATGAGTGATCACCTCATCATTCAATTAT |
| oJZ201 | *GCTGTCTGTCCAAATGGT*AAGCGCATTATC |
| oJZ202 | *ACCATTTGGACAGACAGC*ATCCTTTTTC |
| oJZ212 | TGATCCTGCCTCGCTTGTAC |
| oJZ213 | GACGTCAGGTGGCACTTTTC |
| oJZ214 | TCCGTTGAGAAAGATACTGG |
| oJZ223 | TTCGGGTCGATCATGCCTTG |
| oJZ225 | AACCACTGCTGGGAAACAGG |
| oJZ257 | AGGAGGCTAGCCTATGAATGAGAAAGGATTTACAC |
| oJZ258 | *GAATTCATAAGCTTATACTAGTGTC*TTAATGTTCAACCTTAACTTCTCC |
| oJZ259 | *CATGCATTAGCTAGCATTAC*AGGAGGCTAGCCTATG |
| oJZ394 | *GAAATACATAAAAGATAATATAGGCTGGAA*AGG |
| oJZ395 | *GGAGTTCAAACTTGATATCGAATTCCTG* |
| oJZ396 | *CAGGAATTCGATATCAAGTTTGAACTCC*TTTTTCATATGAGAAG |
| oJZ397 | *TTTTCTGACTCATAGTAGTTCCTCCTT*CTAGATTTTTTTTTGAATTCTACAG |
| oJZ398 | *AAGGAGGAACTACTATGAGTCAGAAAA*CAGACG |
| oJZ399 | *TTCCAGCCTATATTATCTTTTATGTATTTC*TAGCTTCAGAG |
| oJZ436 | CTGGGGGCTGGGATACTTTG |
| oJZ437 | TCTCCCAAGCCAGAACCAAC |

* Restriction sites underlined, Gibson assembly or overlap extension PCR homology flanks italicized

**Supplemental references.**


1. Grant SGN, Jessee J, Bloom FR, Hanahan D. 1990. Differential plasmid rescue from transgenic mouse DNAs into Escherichia coli methylation-restriction mutants. Proc Natl Acad Sci U S A 87:4645–4649.
2. Youngman PJ, Perkins JB, Losick R. 1983. Genetic transposition and insertional mutagenesis in Bacillus subtilis with Streptococcus faecalis transposon Tn917. Proc Natl Acad Sci U S A 80:2305–2309.
3. Chilton SS, Falbel TG, Hromada S, Burton BM. 2017. A conserved metal binding motif in the Bacillus subtilis competence protein ComFA enhances transformation. J Bacteriol 199:e00272-017.
4. Sambrook J, Russell D. Molecular Cloning: A Laboratory Manual Third Edition.
5. Comella BS, Grossman AD, Comella N, Grossman AD. 2005. Conservation of genes and processes controlled by the quorum response in bacteria: characterization of genes controlled by the quorum-sensing transcription factor ComA in Bacillus subtilis. Mol Microbiol 57:1159–1174.
6. Hilgarth RS, Lanigan TM. 2020. Optimization of overlap extension PCR for efficient transgene construction. MethodsX 7:100759.
7. Popp PF, Dotzler M, Radeck J, Bartels J, Mascher T. 2017. The Bacillus BioBrick Box 2.0: expanding the genetic toolbox for the standardized work with Bacillus subtilis. Scientific Reports 2017 7:1 7:1–13.
8. Lemon KP, Grossman AD. 1998. Localization of bacterial DNA polymerase: evidence for a factory model of replication. Science 282:1516–1519.
9. Kraus A, Hueck C, Gartner D, Hillen W. 1994. Catabolite repression of the Bacillus subtilis xyl operon involves a cis element functional in the context of an unrelated sequence, and glucose exerts additional xylR-dependent repression. J Bacteriol 176:1738.
